# *Klebsiella pneumoniae* T6SS exacerbates gut inflammation promoting tumorigenesis

**DOI:** 10.64898/2026.03.31.715367

**Authors:** Konstantinos Fragkoulis, Meri IE Uusi-Mäkelä, Gema Sanz, Cecilia Williams, Ina Schuppe-Koistinen, Ulf O Gustafsson, Lars Engstrand, Staffan Normark, Birgitta Henriques-Normark, Juliette Hayer, Sylvain Peuget, Marie-Stéphanie Aschtgen

**Author notes:** Corresponding authors: Sylvain Peuget, Marie-Stéphanie Aschtgen. co-last authors (Sylvain Peuget, Marie-Stéphanie Aschtgen).

## Abstract

Dysbiosis and bacterial pathobionts contribute to inflammation in IBD and CRC, yet the molecular drivers of this process remain unclear. We identify the *Klebsiella pneumoniae* type VI secretion system (T6SS) as a key promoter of intestinal inflammation and tumor progression. Metagenomic analyses revealed enrichment of T6SS encoding genes in the gut microbiota of IBD patients during inflammatory flares. In zebrafish and mouse models, *K. pneumoniae* T6SS activity exacerbated inflammation and promoted colorectal tumor growth. Mechanistically, T6SS firing enhanced the secretion of LPS via outer membrane vesicles (OMVs), driving NF-κB activation and interferon signalling in host cells. *In vivo*, T6SS-dependent inflammation was associated with the expansion of regulatory T-cell subsets and an immunosuppressive tumor microenvironment. These findings redefine the T6SS as a microbial determinant of host inflammation and cancer progression, highlighting T6SS inhibition as a potential therapeutic approach for IBD and CRC.

**Highlights:**

- T6SS-encoding Enterobacteria are enriched in the gut microbiota of IBD patients
- Klebsiella pneumoniae T6SS exacerbates colitis in mice
- T6SS activity enhances outer membrane vesicle secretion and LPS release
- T6SS promotes colorectal tumorigenesis and immune dysregulation

**Graphical abstract:** 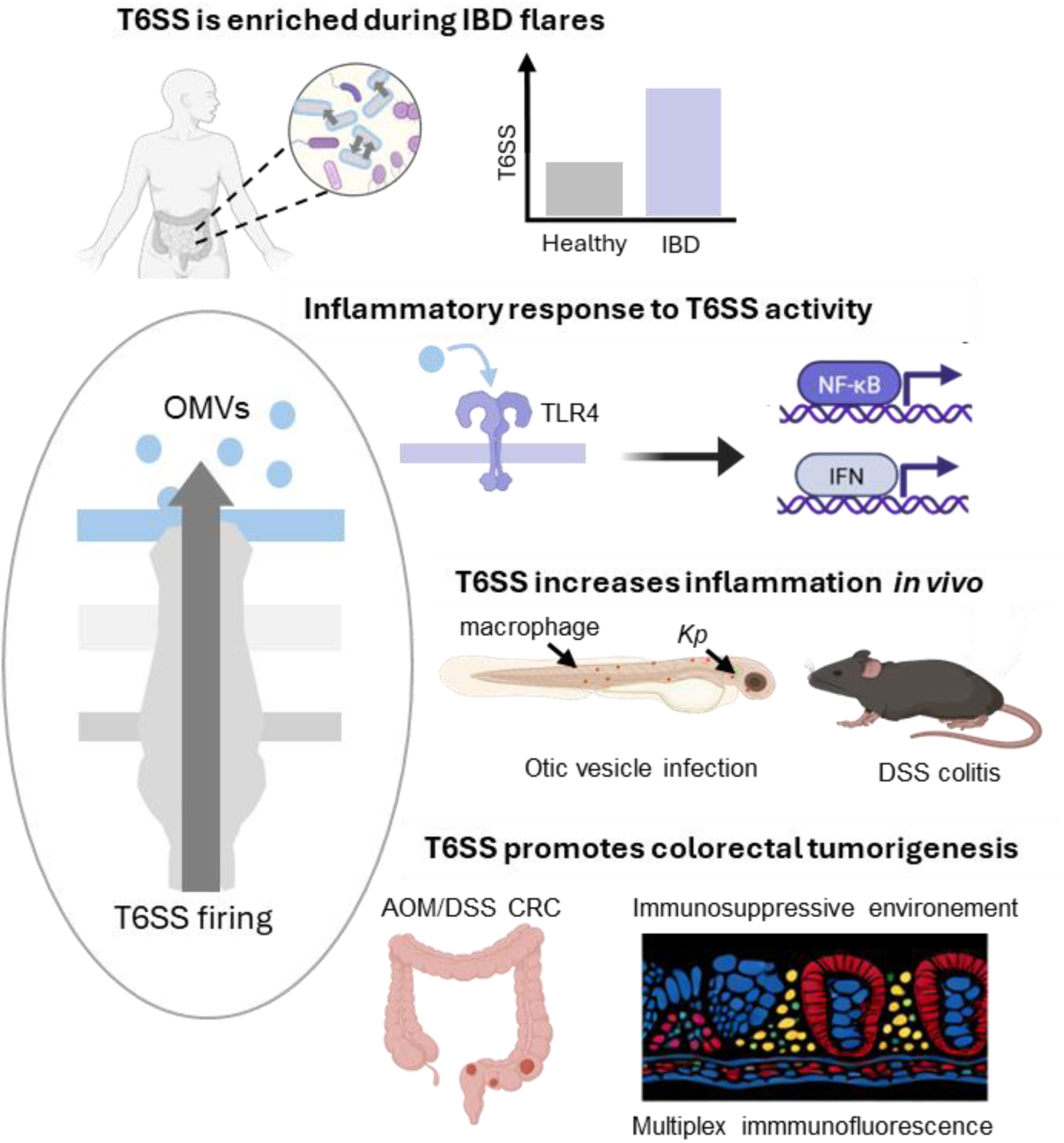

## Introduction

Colorectal cancer (CRC) is a major global health concern, with an increased incidence in patients with inflammatory bowel disease (IBD), where chronic intestinal inflammation drives the development of a pro-tumorigenic microenvironment. The gut microbiota plays a critical role in maintaining intestinal homeostasis, and accumulating evidence suggests that dysbiosis promotes chronic inflammation, thereby contributing to the pathogenesis of both IBD and CRC^1^. Notably, the transfer of IBD-associated dysbiosis microbiota into healthy mice induces intestinal inflammation and accelerates colorectal tumorigenesis^2,3^. Specific pathobionts, including adherent-invasive *Escherichia coli* (AIEC) and *Klebsiella pneumoniae*, have been implicated in shaping dysbiosis, exacerbating intestinal inflammation in IBD, and contributing to tumor development^4–6^.

*K. pneumoniae* has recently emerged as a key pathobiont associated with both IBD exacerbation and CRC development. Metagenomic analyses reveal that *K. pneumoniae* strains are enriched in the gut microbiota of CRC patients^7^. Consistently, in murine models, colonization with *K. pneumoniae* exacerbates colitis and accelerates colorectal tumor growth^8–10^. Mechanistically, we previously showed that *K. pneumoniae* lipopolysaccharide (LPS)-induced inflammation modulates the host p53 tumor suppressor pathway, impacting cancer progression^11^. Moreover, elevated circulating LPS levels and increased TLR4 expression in intestinal tissues are also observed in IBD and CRC patients^12^. Conversely, strategies to inhibit TLR4 signaling have been shown to reduce tumor-promoting inflammation and subsequent tumor burden^13–16^. Despite these insights, the precise mechanisms by which *K. pneumoniae* strains drive inflammation and contribute to colitis and CRC remain poorly understood.

The type VI secretion system (T6SS) is a contractile nanomachine widely used by Enterobacteria to compete for ecological niches and nutrients within the high-density intestinal microbiota. Beyond its role in shaping bacterial interactions, T6SS activity can strongly influence microbial community structure and intestinal homeostasis^17^. In enteric pathogens such as *Salmonella Typhimurium* and *Vibrio cholerae*, T6SS activity reshapes the microbiota composition through interbacterial antagonism and has been linked to heightened inflammatory responses in the host^18,19^. These observations raise the question of how T6SS activity during long-term colonization contributes to intestinal chronic inflammation in IBD and CRC patients.

The T6SS is a contractile, phage-like system that delivers effector proteins into target cells^20^. Unlike other secretion systems, it does not require specific receptors, allowing broad-range targeting of bacteria or host cells. Additionally, the T6SS lacks a defined porin in the outer membrane for pilus passage. Instead, its spear-like structure is thought to locally disrupt the outer membrane, facilitating effector secretion. Recent work also suggests that T6SS effectors can interface with outer membrane vesicles (OMVs), hinting at broader connections between T6SS activity and OMV biology beyond direct antagonism^21^. Here, we propose a novel mechanism by which *K. pneumoniae* T6SS activity promotes gut inflammation through increased OMVs release and associated LPS secretion in the gut microenvironment. By elucidating the role of T6SS in colitis and CRC, our findings provide new insights into *K. pneumoniae*-driven colorectal tumorigenesis and open potential therapeutic strategies targeting bacterial virulence in patients with chronic intestinal inflammation.

## Results

### Enterobacteria T6SS genes are enriched in the gut microbiota of IBD patients

Virulence factor genes (VFGs) differ markedly between commensal and pathogenic Enterobacteriaceae. Recent genomic profiling of *K. pneumoniae* and *E. coli* isolates showed that strains from healthy gut microbiota harbor fewer pathogen-associated VFGs and, notably, lack T6SS effector genes, in contrast to pathogenic isolates that frequently encode complete T6SS effector sets^22^. This suggests that acquisition or enrichment of T6SS-associated functions may be a feature of pathogenic or dysbiosis-associated strains.

To determine whether the gut microbiota of IBD patients is enriched in T6SS-expressing bacteria, we performed metagenomic analyses of the French–Israeli cohort published by Federici *et al*. (Fig. S1A) and of the ulcerative colitis patients from the Swedish KolBiBakt cohort.^6,23^ (Fig. S1B–C). Using a T6SS genomic signature defined by four highly conserved core genes (*tssA, tssB, tssK,* and *tssL*) (Fig. 1A), we observed a consistent enrichment of T6SS Enterobacteriaceae encoding genes in stool samples from both ulcerative colitis (UC) and Crohn’s disease (CD) patients compared with healthy controls, with the highest abundance detected during inflammatory flare episodes (Fig. 1B; Fig. S1D).

**Figure 1.**
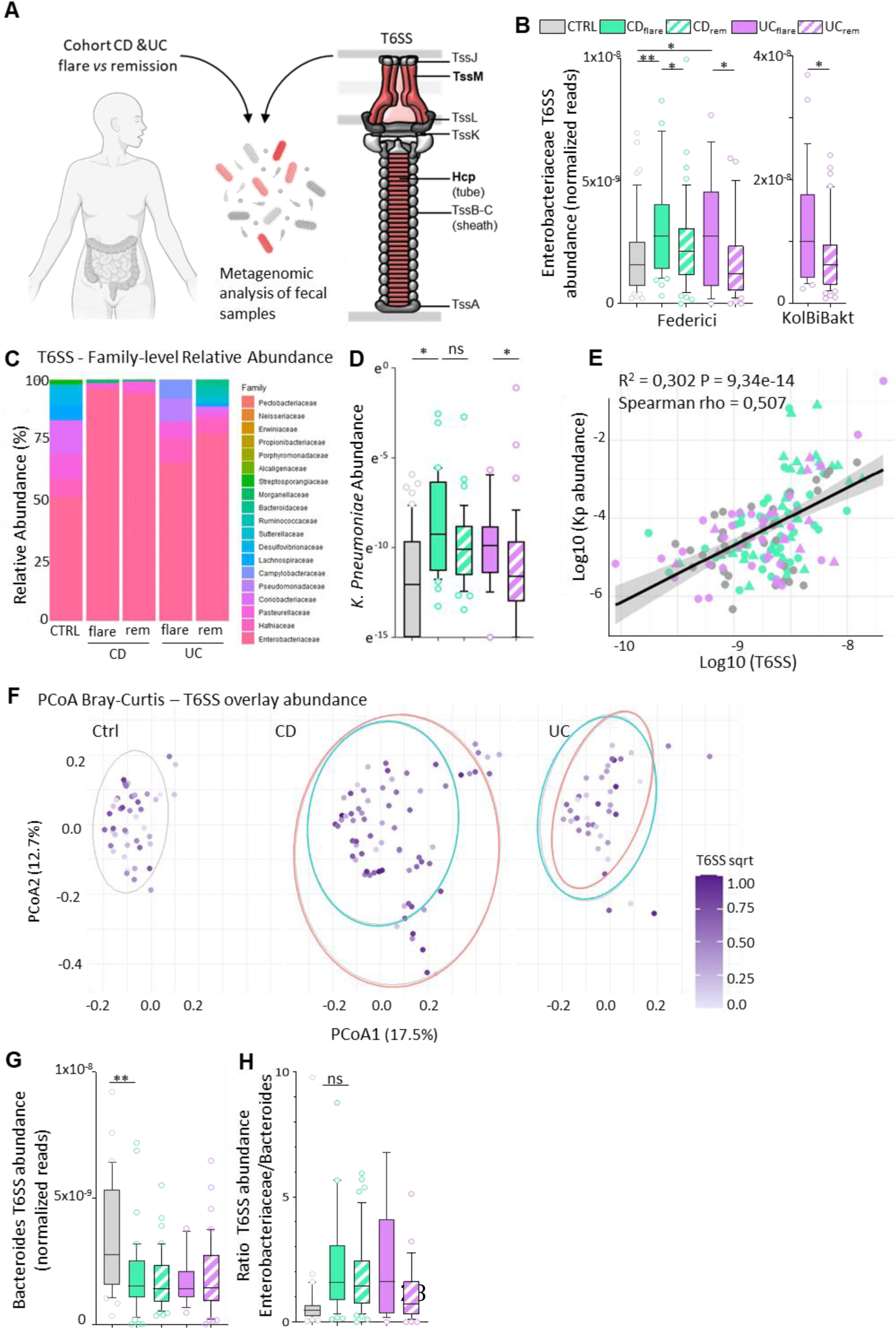
Enterobacteria T6SS genes are enriched in the gut microbiota of IBD patients. (A) Schematic overview of the metagenomic workflow and the Type VI secretion system (T6SS). (B) Normalized abundance of Enterobacteriaceae T6SS core genes (*tssA, tssB, tssK,* and *tssL*) in the Federici cohort (Crohn’s disease and ulcerative colitis) and in ulcerative colitis patients from the KolBiBakt cohort. (C) Family-level relative abundance of taxa contributing to the T6SS signature. (D) Normalized fecal abundance of *Klebsiella pneumoniae* across all samples. (E) Spearman correlation between *K. pneumoniae* abundance and total T6SS gene abundance across all samples. (F) PCoA based on Bray-Curtis dissimilarities, stratified by disease and disease state, colored according to determined T6SS abundance. Ellipses indicate remission (blue) and flare (orange). (G) Normalized abundance of Bacteroides-associated T6SS genes (averaged *tssB*, *tssK*). (H) Ratio of *Bacteroides* to Enterobacteriaceae T6SS signatures. Data are shown as mean ± SD with individual data points overlaid. * p < 0.05; ** p < 0.01.

Taxonomic profiling of T6SS-associated species attributed this enrichment primarily to Enterobacteriaceae. In the French and Israeli cohorts, samples with high T6SS abundance exhibited reduced Shannon diversity, indicating that the signal was dominated by a limited number of taxa rather than broadly distributed across the community (Fig. 1C; Fig S1E). Across the full dataset (Fig. S1F-H), species-level analysis identified *K. pneumoniae* as the most enriched species during flares (Fig. 1D; Fig. S1F-G). Notably, *K. pneumoniae* abundance positively correlated with T6SS gene counts (Fig. 1E). In both CD and UC patients, the microbial community differs between disease states and flare samples correlate with an enriched T6SS signature abundance (Fig. 1F).

Because disease flares are typically associated with depletion of *Bacteroides* spp. and an increased Enterobacteriaceae-to-*Bacteroides* ratio, and given that *Bacteroides* also encode T6SS loci, we next examined lineage-specific T6SS signatures. In contrast to enterobacterial T6SS genes, the *Bacteroides*-associated T6SS signature was enriched in healthy individuals and reduced in IBD patients (Fig. 1G). This pattern paralleled a compositional shift toward Enterobacteriaceae dominance and away from *Bacteroides*, particularly during inflammatory flares (Fig. 1H).

### *K. pneumoniae* T6SS exacerbates inflammation *in vivo*

We first used a zebrafish model to investigate the contribution of *K. pneumoniae* T6SS to inflammation using the innate immune response to the infection as a proxy. GFP-labeled wild-type (WT; non-encapsulated SGH4 T6SS-proficient strain) *K. pneumoniae* and T6SS-deficient isogenic mutants (Δ*hcp* and Δ*tssM*; non-encapsulated SGH4 derivatives) were injected in the otic vesicle of 3 dpf zebrafish larvae harboring mCherry-labeled macrophages [Tg(mpeg1:gal4-UAS:NTR-mCherry) zebrafish]. Then, we performed simultaneous time-lapse imaging of bacterial colonization and macrophage recruitment (Fig. 2A-B; Movies S1-S3). To avoid capsule-mediated immune evasion and systemic infection observed using *K. pneumoniae* SGH4 (encapsulated reference strain), we used non-encapsulated strains with similar growth kinetics for subsequent experiments (Fig. S2A-C). Importantly, the otic vesicle is a sterile environment in which T6SS mutants displayed no colonization defect in absence of bacterial competition, as indicated by comparable GFP fluorescence over time (Fig. S2C). Both WT and T6SS mutants colonized the otic vesicle similarly during the first six hours post-infection and were cleared over the next ten hours. However, WT infection markedly increased macrophage recruitment, as indicated by higher mCherry signal intensity, despite similar bacterial burden (Fig. 2C-D; Fig. S2B-D).

**Figure 2.**
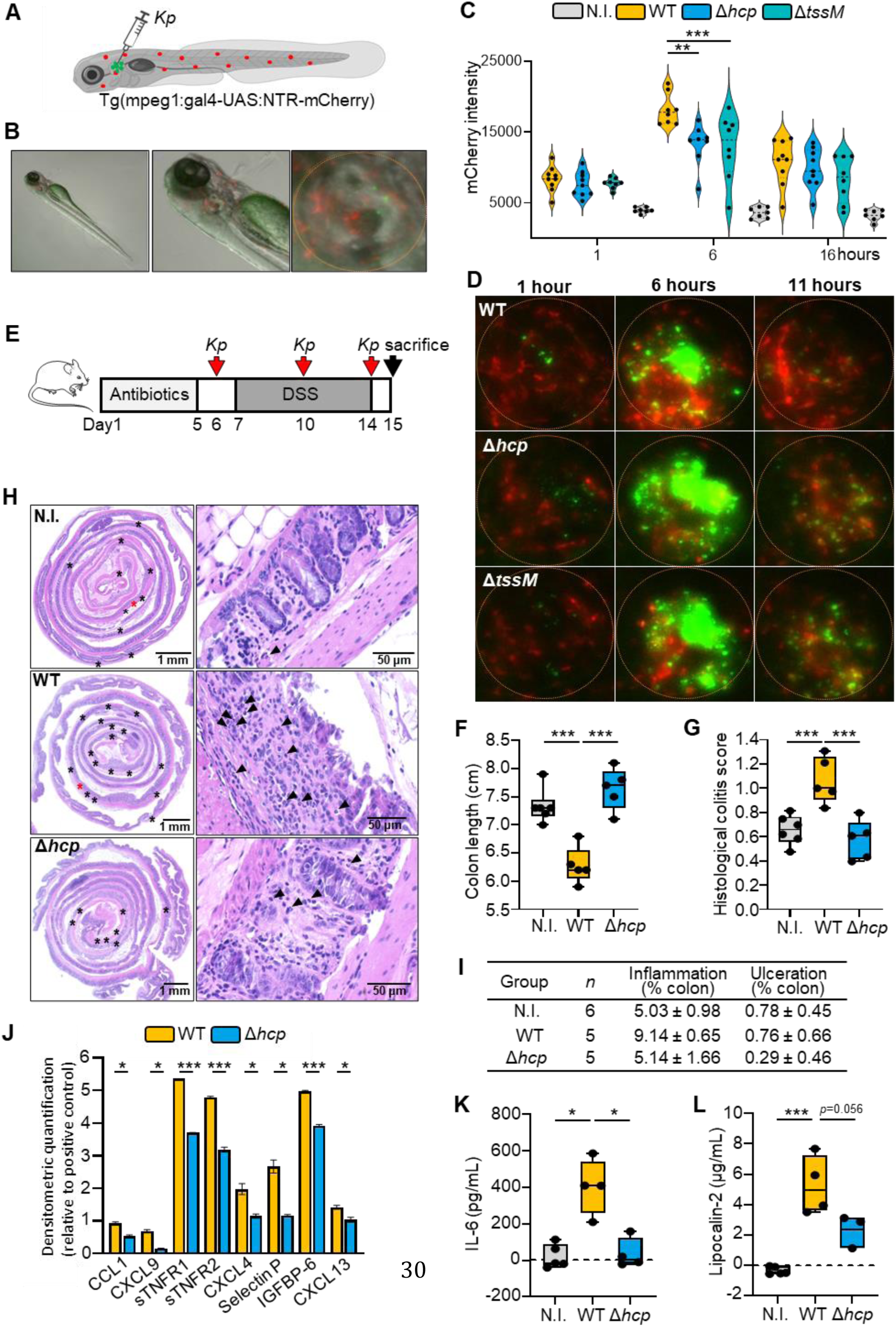
*K. pneumoniae* T6SS activity exacerbates inflammation *in vivo*. (A) Schematic representation of *Tg*(mpeg1:gal4-UAS:NTR-mCherry) zebrafish larvae expressing mCherry-labeled macrophages, microinjected with GFP-labeled *K. pneumoniae* strains into the otic vesicle. (B) Representative overlay image showing brightfield, mCherry macrophages, and GFP-labeled bacteria; the otic vesicle is outlined. (C) Quantification of macrophage infiltration (mCherry fluorescence) in the otic vesicle over time, comparing non-infected larvae (N.I), infected with WT (non-encapsulated SGH4 T6SS-proficient strain) or T6SS-deficient mutant strains (Δ*hcp* and Δ*tssM*; non-encapsulated SGH4 derivatives). ** p < 0.01; *** p < 0.001. Each dot represents an individual zebrafish. (D) Representative images of otic vesicles at indicated time points showing macrophage recruitment and bacterial accumulation. (E) Experimental design for the murine colitis model. Antibiotic-treated C57BL/6 mice were mono-colonized with WT or Δ*hcp K. pneumoniae* by oral gavage, followed by induction of colitis using 2% DSS in drinking water. (F–L) Assessment of colitis severity. (F) Colon length at sacrifice. (G) Histological colitis scores from H&E-stained colon Swiss rolls. (H) Representative H&E-stained sections; stars indicate inflamed (black) or ulcerated (red) regions, and arrows denote infiltrating neutrophils. (I) Criteria used for histological scoring. (J) Relative cytokine levels in serum of WT-and Δ*hcp*-colonized mice at sacrifice, measured by cytokine array. (K–L) IL-6 and lipocalin-2 levels in serum at sacrifice, measured by ELISA. Error bars ± SD. * p < 0.05; *** p < 0.001. Each dot represents an individual mouse.

To further evaluate the contribution of *K. pneumoniae* T6SS to intestinal inflammation, we used an acute dextran sulfate sodium (DSS)-induced colitis model in mice (Fig. 2E). Antibiotic-treated mice were mono-colonized with WT or Δ*hcp* strains before exposure to DSS. We used non-encapsulated strains to reduce virulence while maintaining stable colonization (Fig. S3A)^24^, as in the zebrafish experiments. WT-colonized mice experienced more severe colitis compared to Δ*hcp*-colonized or non-infected controls, as indicated by hallmark features of colonic inflammation, including reduced colon length, higher histological colitis scores, extensive mucosal damage and immune cell infiltration, as well as greater weight loss (Fig. 2F-I; Fig. S3B). In contrast, Δ*hcp*-colonized mice exhibited tissue morphology similar to the controls (Fig. 2H). Consistent with these observations, we detected elevated levels of pro-inflammatory cytokines, including IL-6 and lipocalin-2, in the serum of WT-colonized mice, associated with increased NF-κB activation in the colonic epithelium (Fig. 2J-L, Fig. S2C-G). Together, these results indicate that *K. pneumoniae* T6SS exacerbates host inflammatory response *in vivo* in zebrafish and during colitis in mice.

### *K. pneumoniae* T6SS enhances inflammatory signaling in host cells

T6SS activity is known to mediate interbacterial competition in the intestine, with several enterobacterial T6SS clusters induced under gut-like or high-cell-density conditions^25^. *K. pneumoniae* T6SS expression is generally repressed under standard laboratory conditions such as LB broth but activated under infection-like or competitive environments^26,27^. To confirm this in our experimental settings, we monitored T6SS expression in *K. pneumoniae* SGH4 using a transcription reporter, which revealed T6SS expression only under infection-mimicking and competition conditions (Fig. S4A-B). Functional assessment of T6SS assembly and activity by Hcp secretion assay using FLAG-tagged Hcp strains demonstrated secretion of Hcp exclusively by WT, but not by mutant strains (Fig. S4C). Additionally, we observed comparable growth rates for all strains (Fig. S4D).

Having defined the conditions under which the T6SS is active, we next examined how T6SS activity influences host inflammatory signaling. To assess this, we performed RNA-sequencing on THP-1 macrophages infected with WT or Δ*tssM* strains. WT infection triggered extensive gene expression changes relative to non-infected controls (Fig. S5A), whereas Δ*tssM* infection induced a markedly attenuated response (Fig. S5B). Notably, Gene Ontology (GO) pathway analysis of differentially expressed genes (DEGs) revealed reduced activation of key immune and inflammatory pathways in Δ*tssM*-infected cells compared with WT (Fig. 3A-B). Consistently, gene set enrichment analysis (GSEA) revealed that interferon and inflammatory response were among the most downregulated pathways in Δ*tssM*-compared to WT-infected cell, reflecting a dampened activation of these pathways in absence of T6SS (Fig. 3C-D; Fig. S5C). Transcription factor binding motif analysis of DEG promoters revealed significant enrichment for the Interferon Regulatory Factor (IRF) family and STAT2, in line with activation of the interferon pathway (Fig. S5D).

**Figure 3.**
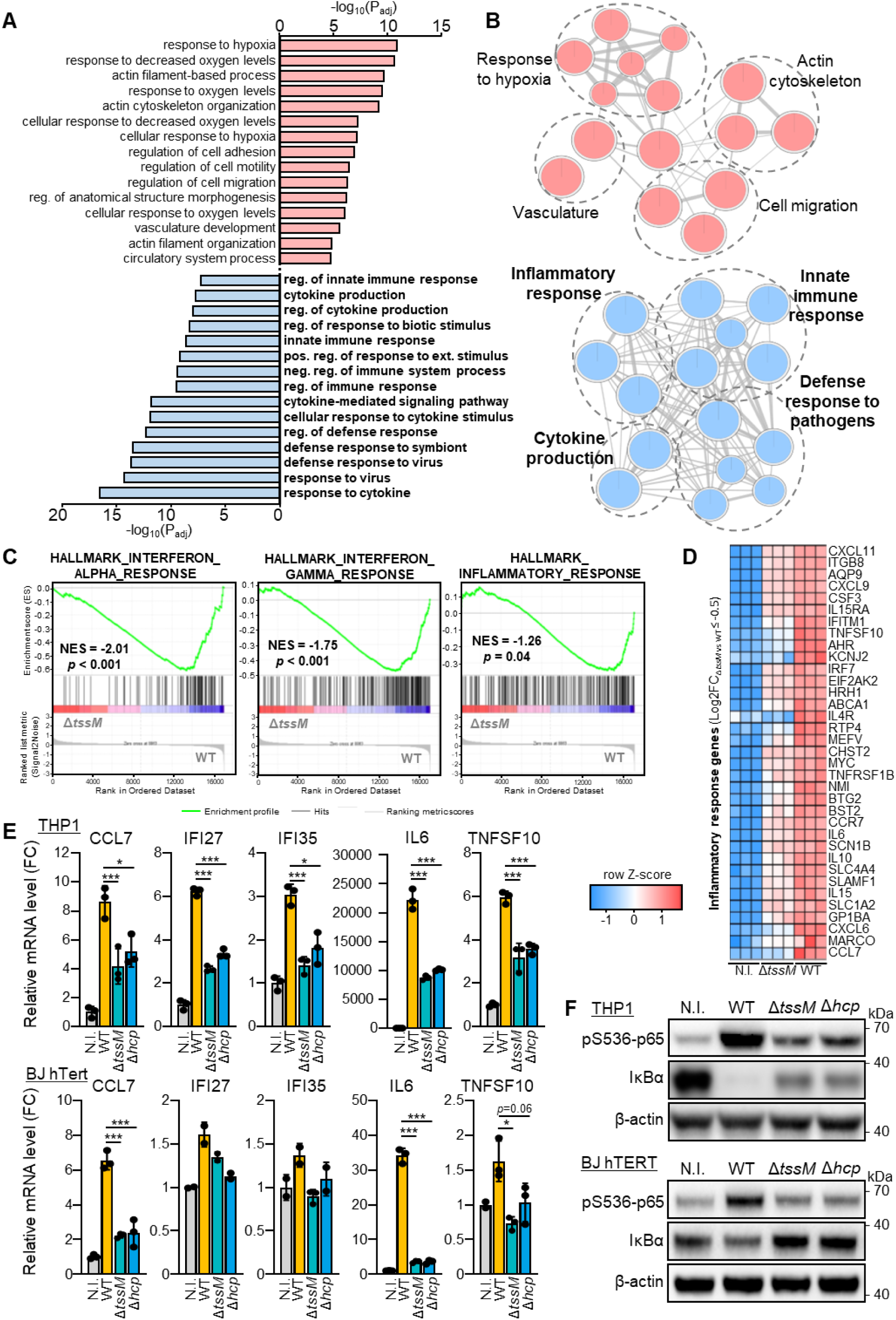
*K. pneumoniae* T6SS induces an inflammatory response and NF-κB activation in infected cells. (A-D) RNA-seq analysis THP-1 macrophages infected with *K. pneumoniae* WT or Δ*tssM* strains: top GO:BP terms enriched in upregulated (red) or downregulated (blue) DEGs in Δ*tssM*-vesus WT-infected cells (A); network visualization of the most significantly up-and downregulated pathways (B); Gene set enrichment analysis (GSEA) comparing Δ*tssM*-vesus WT-infected cells for interferon-α, interferon-γ, and inflammatory response hallmarks (C); Heatmap of DEGs in the inflammatory response (GO:0006954) across conditions (D). N.I.: non-infected. (E) RT-qPCR validation of selected NF-κB target genes in THP-1 and BJ hTERT cells infected with WT, Δ*tssM* or Δ*hcp K. pneumoniae*. Expression is plotted as fold change over non-infected cells. Error bars ± SD. * p < 0.05; *** p < 0.001. (F) Western blots showing NF-κB activation upon infection, as indicated by p65 (Ser536) phosphorylation and IκBα degradation.

To confirm these results, we performed RT-qPCR for inflammatory genes (*IL6*, *CCL7*, *IFI27*, *IFI35* and *TNFSF10*), both in THP-1 macrophages and BJ hTERT fibroblasts infected with *K. pneumoniae* WT, Δ*tssM* or Δ*hcp*. In both cell lines, T6SS mutants showed an average of ∼50% reduction in gene induction compared to WT-infected cells (Fig. 3E). Moreover, we observed impaired NF-κB activation in cells infected with T6SS mutants, as evidenced by reduced p65 Ser536 phosphorylation and decreased IκB degradation (Fig. 3F). In parallel, phosphorylation of TBK1 and IRF7 was markedly reduced, consistent with attenuated activation of the interferon axis (Fig. S5E). Given the role of TBK1 in coordinating NF-κB and IRF signaling, these results suggest that T6SS activity amplifies inflammatory signaling by promoting TBK1-dependent activation of both pathways.

We further confirmed that these findings were independent of the *K. pneumoniae* capsular type by testing encapsulated strains, including SGH4 (serotype K2) and MGH78578 (serotype K52) (Fig. S6A). Moreover, complementation of the T6SS mutants restored NF-κB transcriptional activity to levels comparable with WT-infected cells, as determined by luciferase reporter assays in both THP-1 and BJ hTERT cells (Fig. S6B-C).

Since T6SS can deliver effectors both via contact-dependent and-independent mechanisms, we investigated whether *K. pneumoniae* T6SS-mediated inflammation response required direct bacteria-host contact.

Notably, we found that T6SS-dependent activation of NF-κB and induction of inflammatory genes were recapitulated when bacteria were physically separated from host cells using a permeable transwell system (Fig. 4A), or when cells were exposed to cell-free bacterial supernatants (Fig. 4B-F). Consistent with the infection assays, supernatants from complemented T6SS mutants induced NF-κB at levels comparable with those of WT bacteria (Fig. S7A-B). These findings indicate that *K. pneumoniae* T6SS triggers host inflammatory signaling, at least in part, through a contact-independent mechanism.

**Figure 4.**
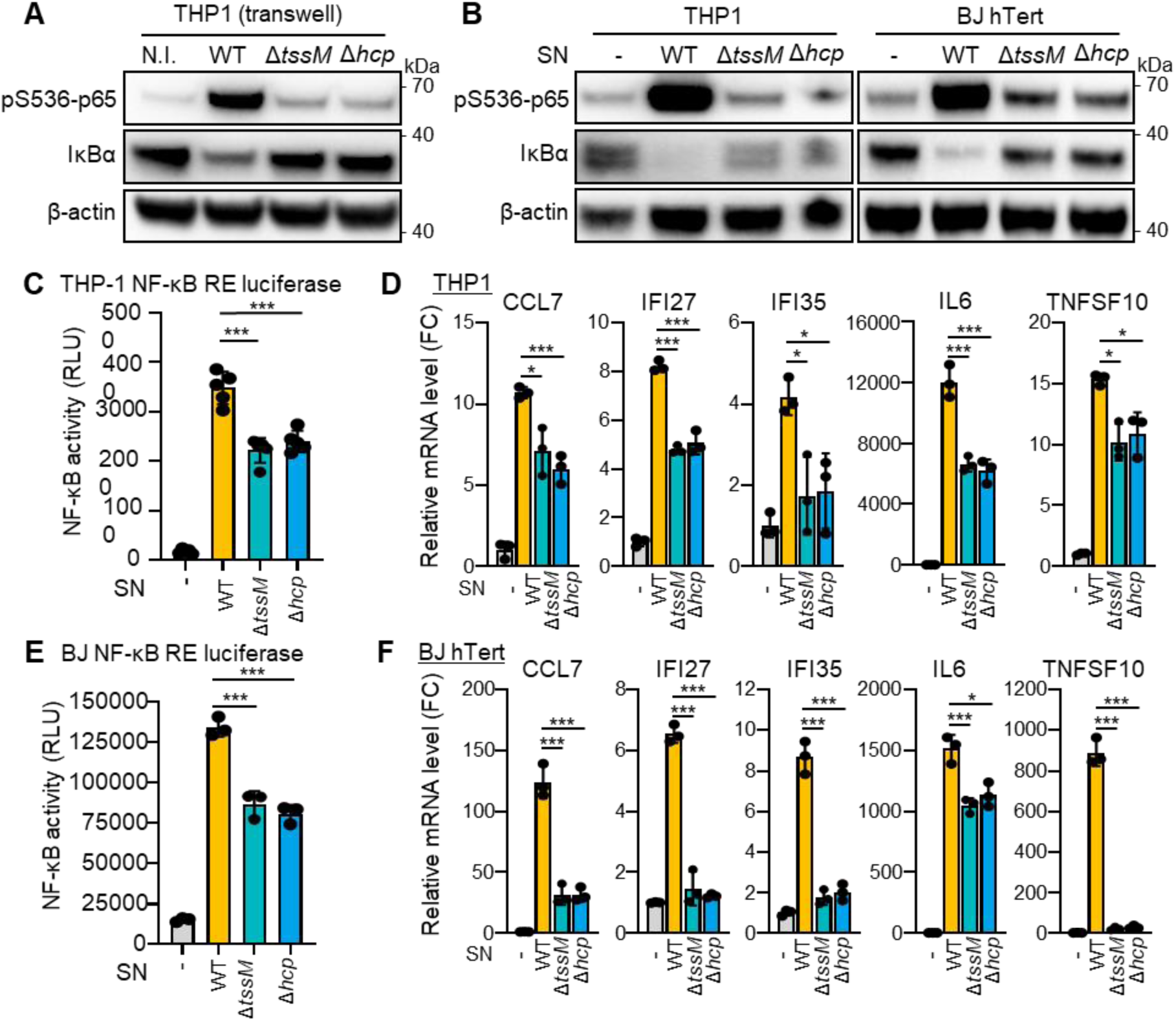
*K*. *pneumoniae* T6SS activates NF-κB through a contact-independent mechanism. (A) Western blot analysis of NF-κB activation, as indicated by p65 (Ser536) phosphorylation and IκBα degradation, in THP-1 cells co-cultured with *K. pneumoniae* WT, Δ*tssM*, or Δ*hcp* strains using a Transwell system to prevent direct bacteria–host contact. (B) Western blots showing NF-κB activation in THP-1 and BJ hTERT cells treated with cell-free bacterial supernatants (SN) from *K. pneumoniae* WT, Δ*tssM*, or Δ*hcp* cultures. (C, E) NF-κB transcriptional activity measured by luciferase reporter assay in THP-1 (C) and BJ hTERT (E) NF-κB reporter cells following treatment with bacterial SN. Error bars ± SD. *** p < 0.001. (D, F) RT-qPCR quantification of selected NF-κB target genes in THP-1 (D) and BJ hTERT (F) cells following treatment with bacterial SN. Error bars ± SD. * p < 0.05; *** p < 0.001.

### *K. pneumoniae* T6SS activity increases OMVs secretion and LPS release

We previously demonstrated that NF-κB activation in fibroblasts by WT *K. pneumoniae* supernatant is largely TLR4-dependent^11^, suggesting that pattern recognition receptor (PRR) signaling could mediate the T6SS-dependent phenotype we observed under similar conditions. Indeed, TLR4 inhibition by TAK-242 or supernatant from Δ*lpxM* strains which produce an antagonist TLR4 LPS form, prevented T6SS-dependent activation of NF-κB in THP1 cells (Fig. 5A). We next examined whether *K. pneumoniae* T6SS modulates the secretion of immune-stimulatory factors and microbe-associated molecular patterns (MAMPs). LPS quantification showed reduced levels in *K. pneumoniae* supernatants from T6SS mutants compared to WT (Fig. 5B), in line with prior publications linking T6SS to MAMPs release^28^. LPS is typically secreted within membrane-bound OMVs^29^, and correspondingly, WT bacteria released significantly more lipids than T6SS mutants, indicating increased OMVs production (Fig. 5C). Importantly, a similar T6SS-dependent increase was observed in *K. pneumoniae* encapsulated strains under T6SS expression condition (Fig. S8A-C), and LPS and lipid secretion were restored in Δ*hcp* and Δ*tssM* complemented strains (Fig. S8D-E). Collectively, these results indicate that T6SS activity promotes LPS release via OMVs, providing a mechanistic link between T6SS and the host inflammatory response.

**Figure 5.**
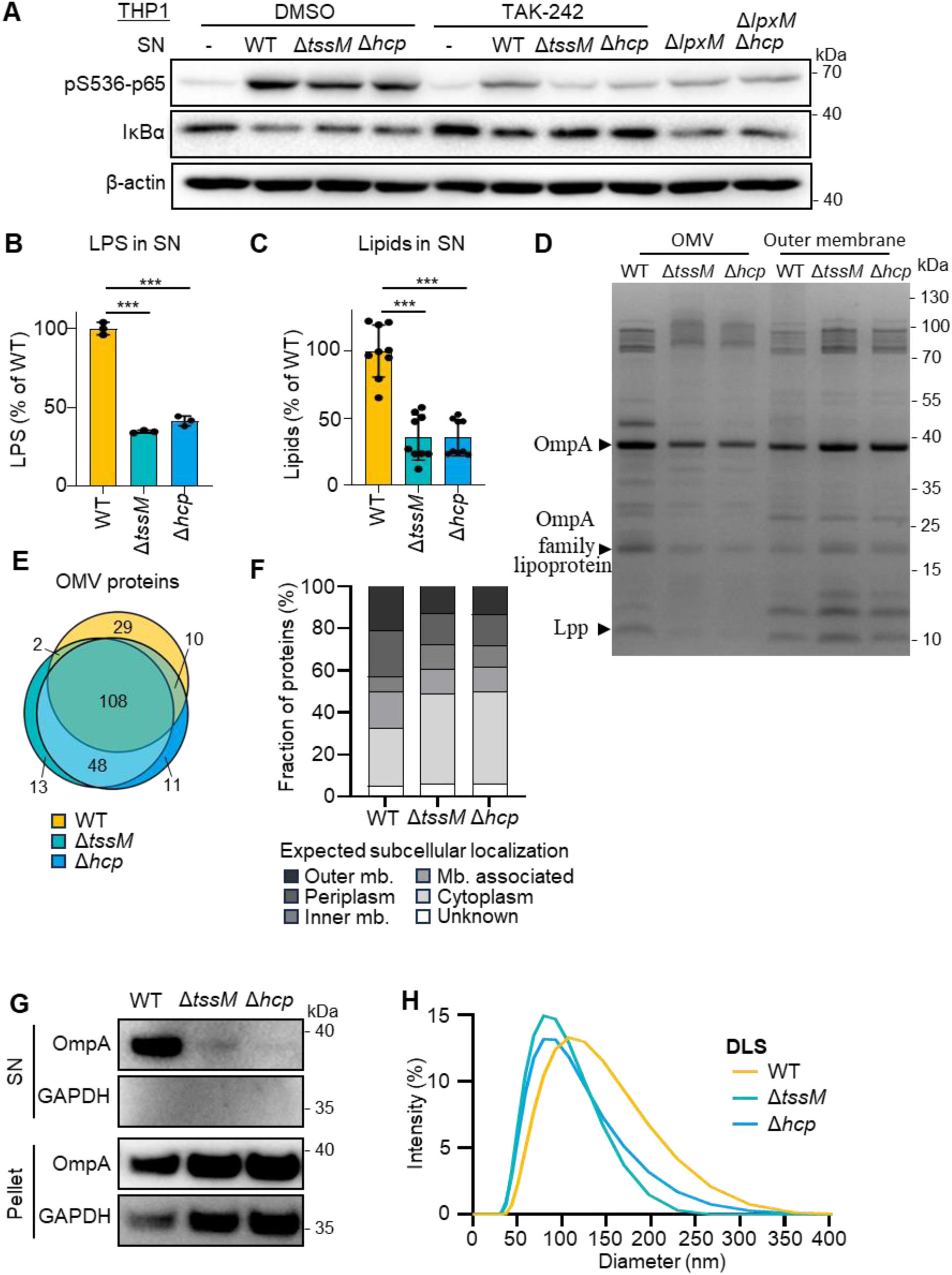
T6SS activity increases OMVs secretion and LPS release. (A) Western blot analysis of NF-κB activation in THP-1 cells, assessed by p65 (Ser536) phosphorylation and IκBα degradation, following treatment with bacterial supernatants (SN) from *K. pneumoniae* WT, Δ*tssM,* Δ*hcp*, Δ*lpxM*, or double mutant Δ*lpxM*Δ*hcp* strains. TLR4 involvement was verified using the TLR4 inhibitor TAK-242. (B) LPS quantification in bacterial SNs by LAL assay. Error bars ± SD. *** p < 0.001. (C) Total lipid content in SNs measured by FM4-64 fluorescence. Error bars ± SD. *** p < 0.001. (D) Comparison of protein profiles between purified OMVs and corresponding outer membrane (OM) fractions by SDS-PAGE and silver staining. Selected protein bands (arrows) were excised and identified by mass spectrometry. (E) Venn diagram showing overlap of proteins identified by mass spectrometry in OMVs from WT and T6SS mutant strains. (F) Predicted subcellular localization of proteins identified in OMVs preparations. (G) Detection of OMVs by Western blot using OmpA as a marker in both SN and pellet fractions. (H) OMVs size distribution analyzed by dynamic light scattering (DLS).

To assess whether T6SS activity influences OMVs composition, we then compared the SDS–PAGE protein profiles of purified OMVs isolated from WT and T6SS mutant strains. Several proteins displayed altered abundance in the absence of T6SS (Fig. 5D). Notably, three major OMV-associated proteins (OmpA, an OmpA-associated lipoprotein, and Lpp) were almost undetectable in OMVs from T6SS mutants. Mass spectrometry confirmed these differences in protein abundance, although the overall OMVs proteome remained largely conserved between strains (Fig. 5E-F; Fig. S9A-B). Consistent with its established association with OMVs^30^, OmpA was the most abundant protein in WT OMVs, and was enriched in WT supernatants compared with T6SS mutants (Fig. 5G). Dynamic light scattering (DLS) analysis further revealed that WT-derived OMVs were slightly larger (average diameter ∼115 nm) than those from T6SS mutants (∼92 nm) (Fig. 5H).

LPS profiling of purified OMVs showed that the lipid A composition was conserved between strains (Fig. S9C). In contrast, the protein profile of WT-derived OMVs closely resembled that of the WT OMs, whereas OMVs from T6SS mutants displayed a distinct protein composition compared to their corresponding OM fractions (Fig. 5D). These findings suggest that the T6SS activity influences the mechanism of OMV biogenesis, possibly by inducing local membrane instability associated with T6SS firing events.

### *K. pneumoniae* T6SS promotes colorectal tumorigenesis and fosters an immunosuppressive microenvironment

To examine the contribution of *K. pneumoniae* T6SS to inflammation-driven colorectal tumorigenesis, we established a mono-colonization model coupled with AOM-DSS treatment (Fig. 6A). Over the course of the experiment, WT-colonized mice exhibited greater weight loss than Δ*hcp*-colonized, despite comparable bacterial colonization (Fig. S10A-B). Moreover, at the experimental endpoint, WT-colonized mice developed higher number and larger size of colonic tumors compared with Δ*hcp*-colonized mice (Fig. 6B-D; Fig. S9C-E), associated with more advanced dysplasia and increased epithelial proliferation, as indicated by Ki-67 staining (Fig. 6E-G). These findings indicate that the *K. pneumoniae* T6SS promotes inflammation-associated colorectal tumor development.

**Figure 6.**
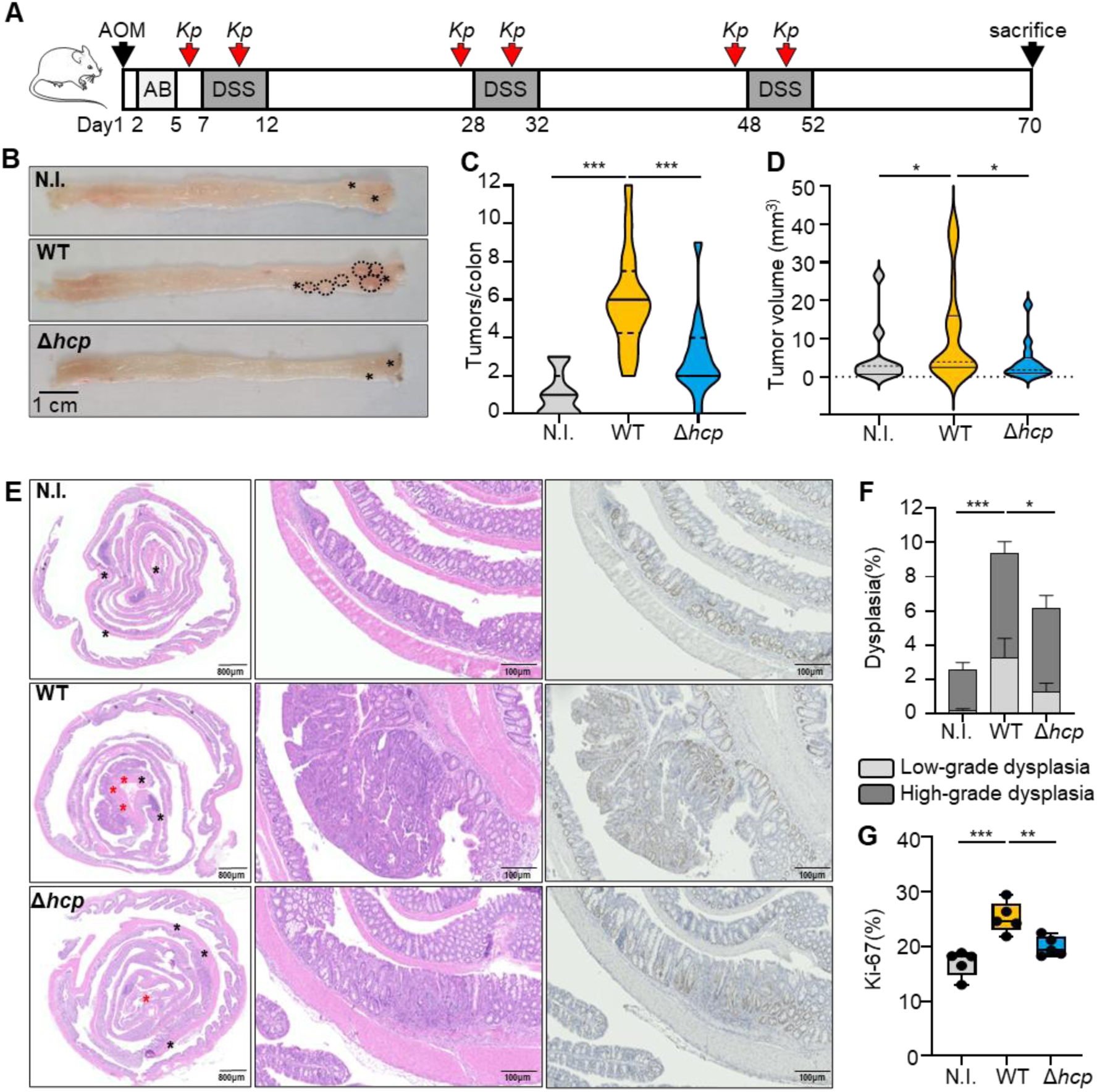
*K. pneumoniae* T6SS promotes inflammation-driven colorectal tumorigenesis. (A) Experimental design of the AOM/DSS experiment to induce colitis-associated colorectal cancer in mice mono-colonized with WT or Δ*hcp K. pneumoniae*. (B) Representative macroscopic images of colons at sacrifice. Small tumors (<4 mm³) are indicated by stars; large tumors (>4 mm³) are outlined with dashed lines. (C) Quantification of visible tumor counts per colon. *** p < 0.001. (D) Distribution of individual tumor volumes across experimental groups. * p < 0.05 (E) Representative histological sections of H&E-stained colon Swiss rolls and Ki-67 immunostaining. Dysplastic areas are indicated with a star (red: high grade, black: low grade). (F) Quantification of the proportion of dysplastic areas per colon in each group. Statistical comparisons were performed on total dysplasia percentages between groups. Error bars ± SD * p < 0.05; *** p < 0.001. (G) Quantification of epithelial cell proliferation via Ki-67-positive cell counts. Error bars ± SD. * p < 0.05; *** p < 0.001.

To define the immune landscape associated with T6SS-dependent tumorigenesis, we performed sequential multiplex immunofluorescence (mIF) using ten markers to profile eight distinct immune cell populations in the whole colon of infected mice (Fig. 7A-B). WT-colonized mice exhibited a pronounced increase in T cells, particularly regulatory subsets. CD3⁺ CD4⁺ FOXP3⁺ regulatory T cells (T4regs) accumulated markedly in WT-colonized mice relative to Δhcp-colonized or non-infected controls (Fig. 7C-D), consistent with their known role in establishing immune tolerance within the tumor microenvironment^31^. We also observed a T6SS-dependent increase in CD3⁺ CD8⁺ FOXP3⁺ regulatory T cells (T8regs), a gut-specific population implicated in immunosuppression and CRC progression^32,33^ (Fig. 7E–F). Both T4regs and T8regs were localized adjacent to tumor sites (Fig. 7C-F), suggesting that *K. pneumoniae* T6SS-driven inflammation is associated with expansion of regulatory T cells and local immunosuppression, thereby fostering a tumor-permissive microenvironment.

**Figure 7.**
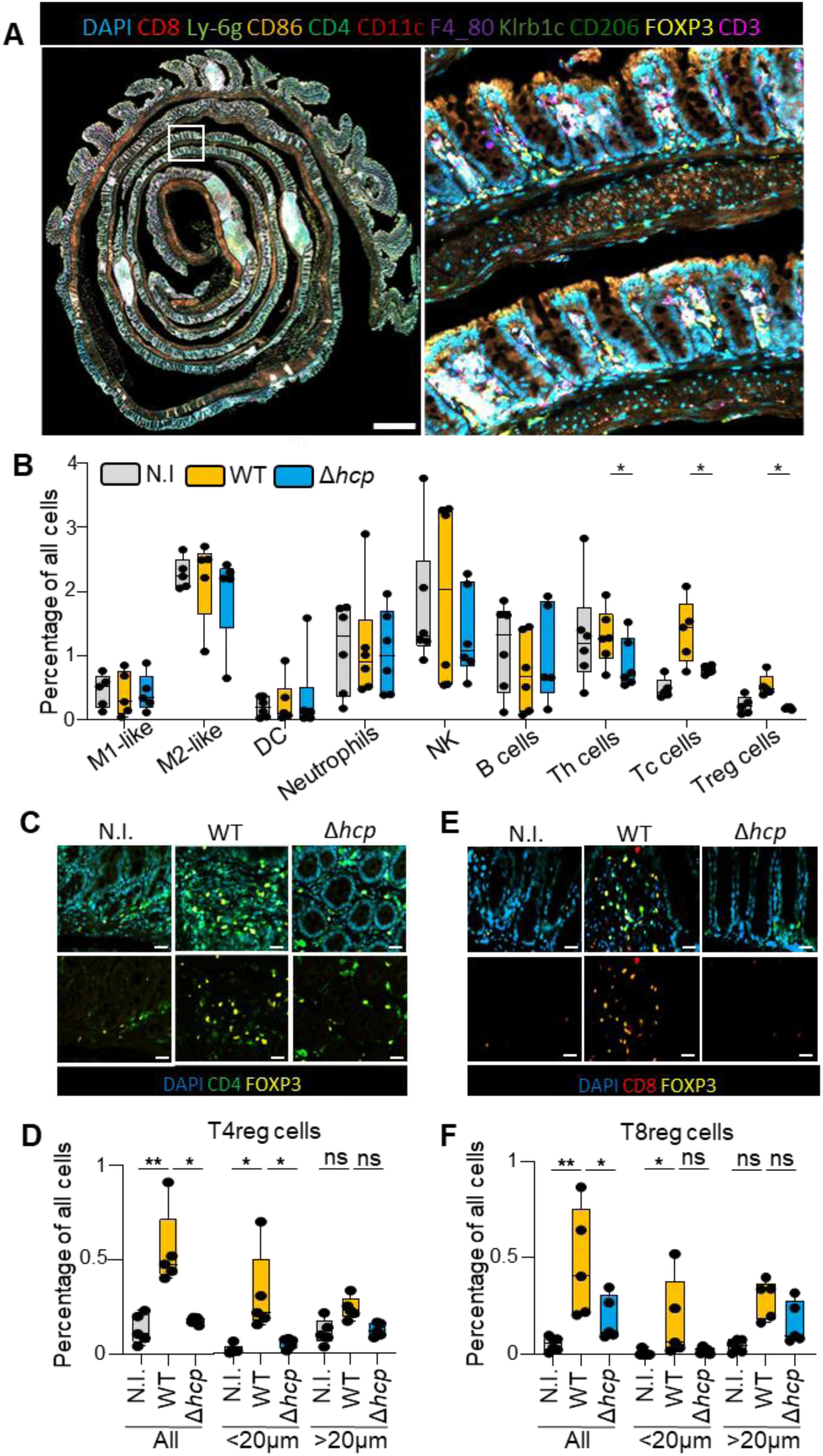
*K. pneumoniae* T6SS activity promotes accumulation of immunosuppressive regulatory T cells in the tumor microenvironment. (A) Multiplex immunofluorescence (mIF) analysis of colon tissues from AOM/DSS-treated mice colonized with WT or Δ*hcp K. pneumoniae*. Ten immune markers were profiled: Ly-6G (lime), CD8 (red), CD3 (magenta), FOXP3 (yellow), CD86 (orange), CD11c (dark red), CD4 (green), Klrb1c (olive), F4/80 (purple), and CD206 (dark green). Scale bars, 200 μm. (B) Quantification of the relative abundance of each immune cell population across experimental groups. Error bars ± SD. * p < 0.05. (C–D) CD4⁺FOXP3⁺ regulatory T cells (T4regs) within colon tissue: representative mIF image (C) and quantification in proximity to tumor regions (D). (E–F) CD8⁺FOXP3⁺ regulatory T cells (T8regs) within colon tissue: representative mIF image (E) and quantification in proximity to tumor regions (F). Error bars ± SD. * p < 0.05; ** p < 0.01; ns not significant.

## Discussion

Metagenomic studies have revealed strong correlations between specific bacterial taxa and intestinal dysbiosis in IBD, where Proteobacteria are consistently enriched. Among these, AIEC and *K. pneumoniae* are associated with Crohn’s disease. However, the molecular mechanisms underlying their contribution to chronic inflammation remain incompletely understood. One important factor in IBD is the interbacterial competition for ecological niches and resources. Intestinal colonization by AIEC, for instance, alters the microbiota composition and causes chronic colitis in mice, with both phenotypes persisting well beyond AIEC clearance^34^. Similarly, *K. pneumoniae* has been associated with flare episodes in IBD patients^6^, underlying the long-lasting influence of bacterial competition on gut homeostasis and inflammation.

The T6SS is a major bacterial weapon for interbacterial competition within the dense gut microbiota. Beyond its well-described function in bacterial competition^35,36^, T6SS effectors can also modulate host responses and promote inflammation^19^. Our study reveals that *K. pneumoniae* T6SS activity exacerbates inflammation and promotes tumorigenesis through the secretion of LPS via OMVs release, adding another layer in the contribution of T6SS to disease progression. T6SS are predominantly found in Proteobacteria^37^, and our metagenomic analysis of IBD cohorts demonstrates a correlation between Proteobacterial enrichment and the abundance of T6SS-encoding species, particularly during disease flares dominated by *E. coli* and *K. pneumoniae*. Hence, our findings reinforce the idea that T6SS activity exacerbates host inflammation in IBD.

Using mono-colonization experiments, we demonstrate that T6SS activity exacerbates colitis independently of microbiota interactions, establishing its direct contribution to host inflammation. *K. pneumoniae* infection induces a robust NF-κB activation *in vitro* and triggers immune cell recruitment *in vivo* in a T6SS-dependent manner, consistent with previous reports linking T6SS to host immune modulation^28^. Building on our earlier work demonstrating that *K. pneumoniae* induces a significant inflammatory response through LPS-associated OMVs via TLR4 signaling in similar cell models^11^, we hypothesized that T6SS could influence OMVs release and composition. Indeed, our experiments confirmed that the T6SS-mediated inflammatory response is driven by secreted OMVs and its associated LPS. Notably, the ability of T6SS effectors to elicit a strong NF-κB response in the absence of direct bacterial-host contact highlights its role in secreting immunomodulatory components, in line with earlier reports showing that T6SSs can facilitate host-bacteria interactions through secreted MAMPs^28,38^.

Mechanistically, the absence of a defined outer membrane porin to facilitate the passage of the T6SS Hcp/VgrG/PAAR needle suggests that, similar to its bacteriophage counterpart, T6SS can transiently perforate the outer membrane through mechanical force. The force generated by the T6SS sheath contraction is sufficient to propel the T6SS through the membrane^39^. We propose that T6SS firing and outer membrane perforation locally destabilizes the membrane and promotes OMVs release. Supporting this model, our proteomic analysis revealed key differences in OMVs composition between wild-type and T6SS mutant strains. Lpp is a lipoprotein that covalently binds the peptidoglycan layer and is known to be selectively depleted in OMVs^40^. Its higher abundance in wild-type OMVs compared to those from T6SS-deficient strains suggest that OMVs production in presence of T6SS could occur through membrane destabilization rather than canonical bulging mechanisms. Considering that T6SS firing events are estimated to occur at rates exceeding 10^9^ events per minute per gram of colonic content^17^, constant T6SS activity could substantially enhance OMVs production in the gut and subsequent inflammation. Although *K. pneumonaie* T6SS expression is induced mainly in host-associated conditions, this makes these effects most apparent *in vivo*. Interestingly, other bacteria appear to exploit this phenomenon: *Cupriavidus necator* secretes a T6SS effector that binds LPS and recruits OMVs^21^, whereas *Helicobacter hepaticus* and *Bacteroides fragilis* T6SS are linked to reduced inflammation *in vivo*, possibly due to the expression of a TLR4 antagonistic LPS^41–43^.

Our findings also establish a link between *K. pneumoniae* T6SS activity and inflammation-driven colorectal tumorigenesis. Consistent with previous reports, gut colonization with *K. pneumoniae* promotes adenoma growth and progression in the AOM-DSS model^9^, and we further demonstrate that this effect is T6SS-dependent. Elevated circulating cytokines levels and increased dysplasia and cell proliferation in the colons of WT-colonized mice indicate that T6SS-induced inflammation promotes tumor progression, consistent with the well-established role of chronic inflammation in colorectal cancer^44^. Moreover, T6SS activity appears to remodel the immune landscape during chronic colonization. A previous elegant single-cell profiling study has demonstrated that *K. pneumoniae* T6SS reshapes the pulmonary immune landscape by promoting less active immune subsets and altering macrophage targeting. Rather than inducing broad inflammatory activation, T6SS activity favored immune states associated with bacterial persistence^45^. Consistent with these findings, in our model, we observed a T6SS-dependent expansion of gut-resident CD3^+^ CD8^+^ Foxp3^+^ regulatory T cells, a subpopulation known to maintain mucosal tolerance in IBD^46^ but also to facilitate immune evasion in tumors^32^. Given that high Treg infiltration correlates with poor CRC prognosis^47^, our data suggest that a microbiota enriched in T6SS-expressing *K. pneumoniae* or Proteobacteria could negatively impact clinical outcomes in cancer patients.

Together, these findings redefine the T6SS as not only a bacterial competition system but also a determinant of host inflammation and cancer progression through OMV-mediated LPS secretion. This discovery bridges bacterial antagonism with host pathophysiology, underscoring the capacity of microbial warfare systems to reshape the gut microenvironment and promote disease. Targeting T6SS activity may therefore offer a novel therapeutic strategy to restore microbial homeostasis, mitigate inflammation, and improve immune response in IBD and CRC patients. Future studies should explore the efficacy of T6SS-targeting compounds in mitigating dysbiosis and reducing CRC risk in susceptible individuals.

## Resource availability

### Lead contact

Further information and requests for resources and reagents should be directed to and will be fulfilled by the lead contact, Marie-Stéphanie Aschtgen.

### Materials availability

Reagents generated in this study are available, upon reasonable requests, with a completed material transfer agreement.

### Data and code availability

The datasets generated in the study are available in GEO database (https://www.ncbi.nlm.nih.gov/geo/) under access number GSE325990.

## Acknowledgments

We thank our colleagues for their support and for generously sharing protocols, particularly Dr. Matilda Holm for COMET settings. We are grateful to Anny Mais, Dimitrios Salgkamis, Evangelos Tzoras, Nikolaos Tsiknakis for technical support and Dr. Stefanie Prast-Nielsen for help with data management. We also acknowledge the Spatial Proteomics unit at SciLifeLab, the Proteomics Facility at Uppsala University for providing access to instrumentation and technical support and the Animal facility at the Karolinska Institutet. The authors acknowledge the ISO 9001 certified IRD i-Trop HPC (South Green Platform) at IRD Montpellier, as well as the Institut Français de Bioinformatique, financed under the Programme d’Investissements d’Avenir funded by the Agence Nationale de la Recherche (RENABI-IFB ANR-11-INBS-0013), for providing HPC resources that have contributed to the research results reported within this paper. This work was supported by grants from the Swedish Research Council, the Knut and Wallenberg Foundation and Torsten Söderberg Foundation to BHN; the Swedish Cancer Society, the Swiss Bridge Foundation, and the Karolinska Institute to SP; and the Gustavsson Funden and the European Crohn’s and Colitis Organization (ECCO) to MSA.

## Author contributions

Conceptualization: MSA, SP, JH, KF; Methodology: MSA, SP, JH, BHN, SN, CW, GS, MUM, KF; Formal analysis: KF, GS, JH, SP, SN, BHN, MSA; Investigation: KF, MUM, JH, SP, MSA; Resources: MSA, SP, JH, BHN, SN, LE, UG, ISK, CW, GS, MUM, KF; Supervision: SP, MSA; JH, BHN; Funding acquisition: BHN, SN, SP, MSA; Writing – original draft: KF, SP, MSA; Writing – review & editing: All authors.

## Declaration of generative AI and AI-assisted technologies in the writing process

We declare that generative AI and AI-assisted technologies were used only for language editing and proofreading of the manuscript. No AI tools were used to interpret data, generate results, or influence the scientific content of this work.

## STAR★Methods

### KEY RESOURCE TABLE

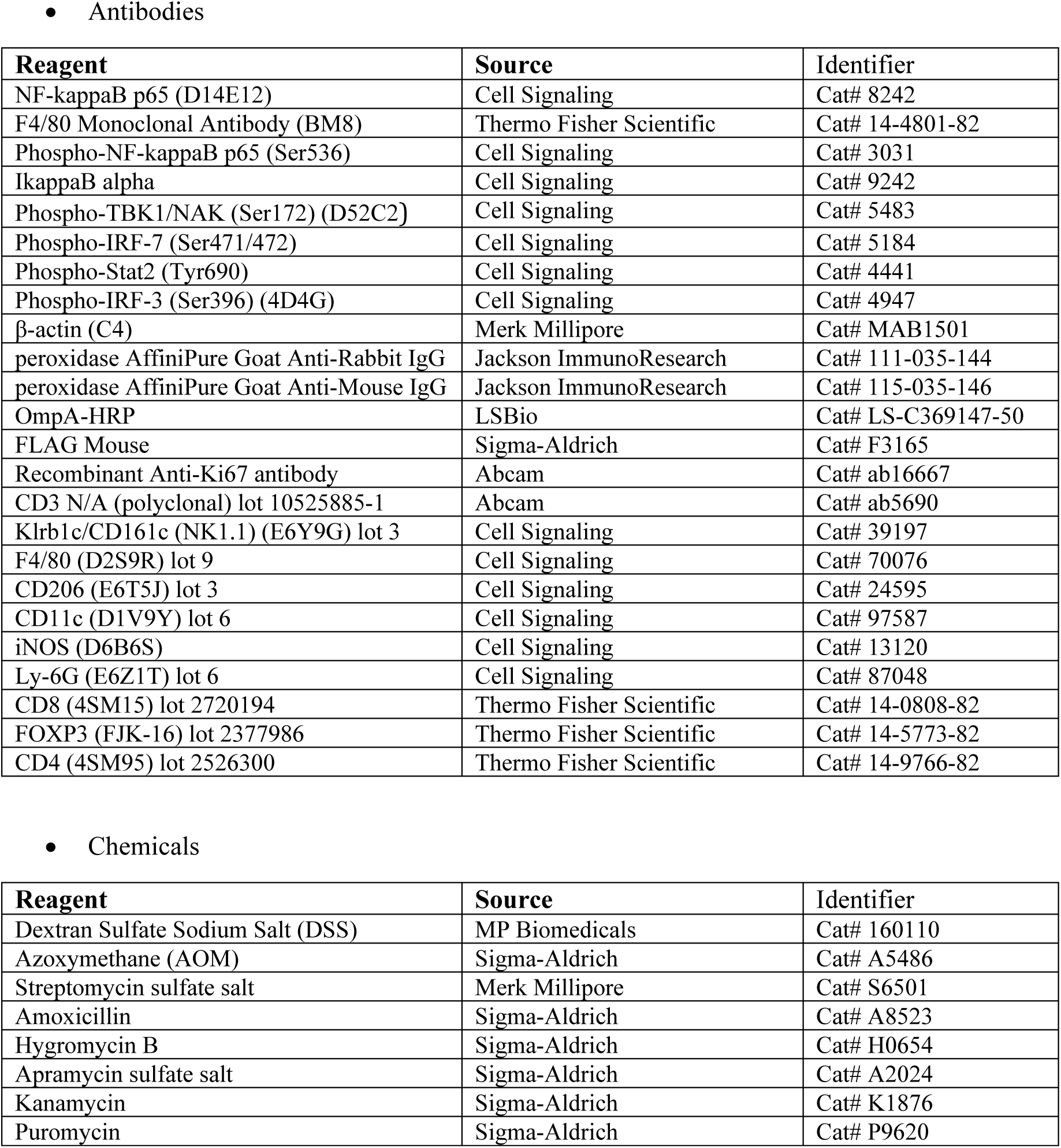
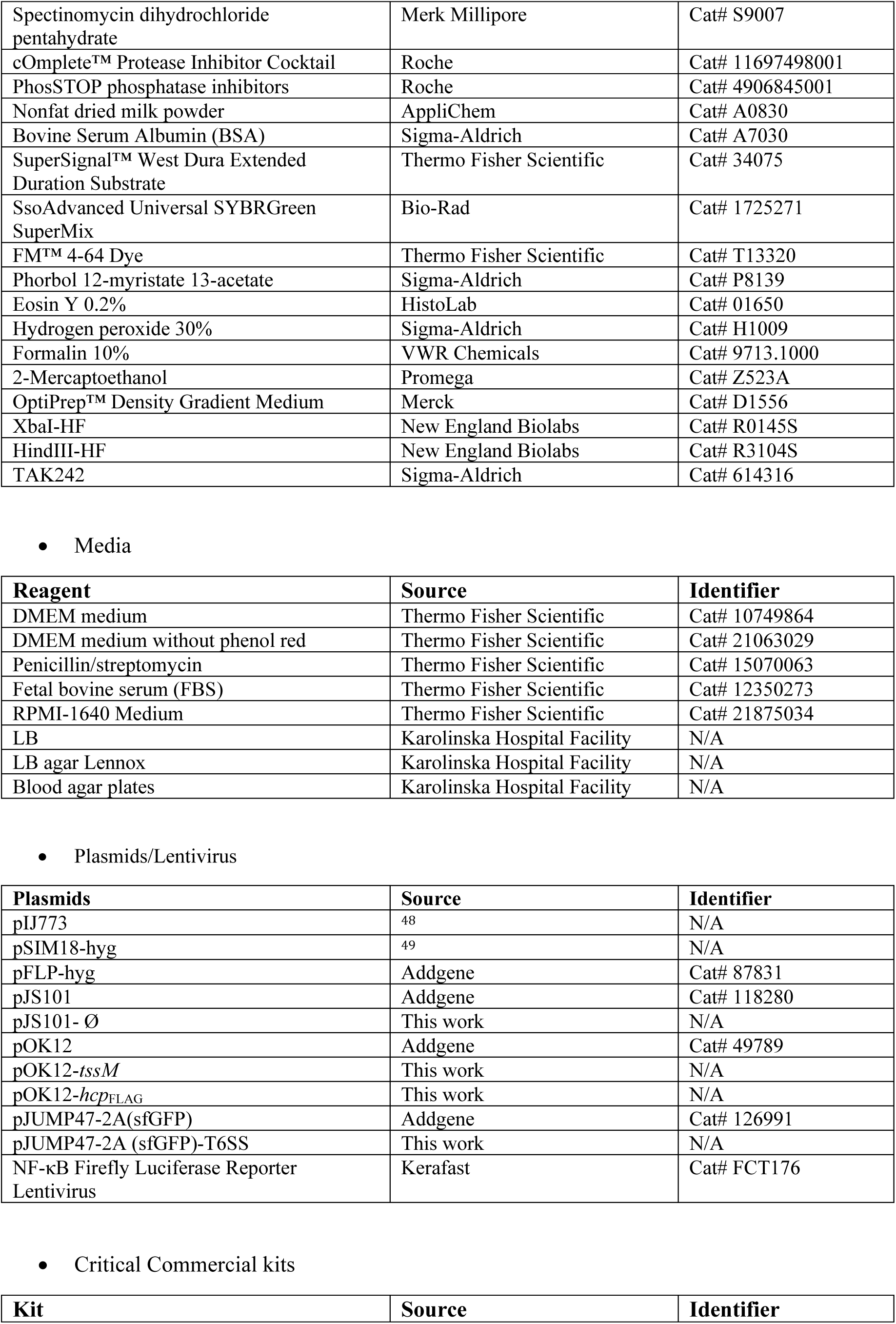
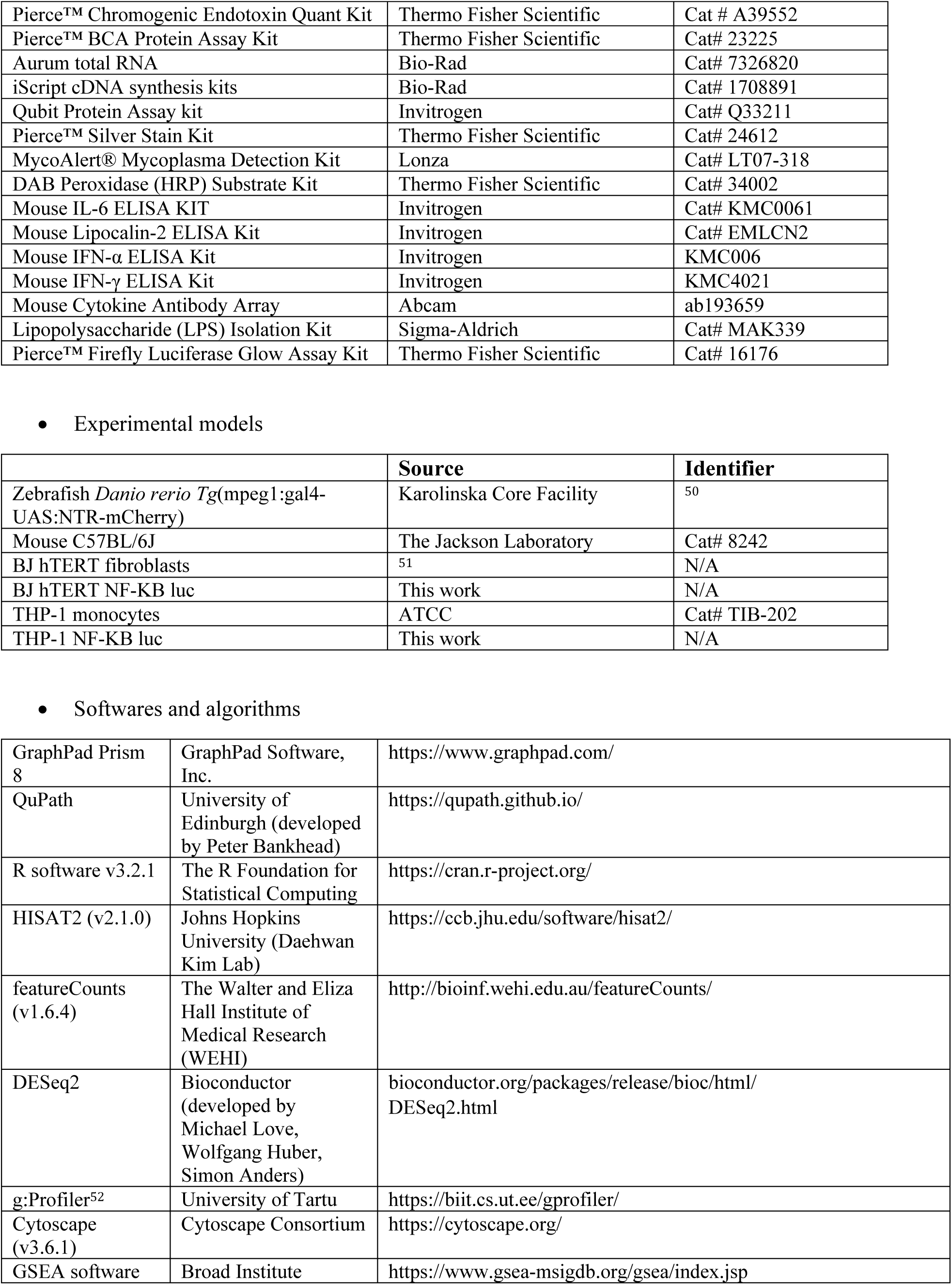
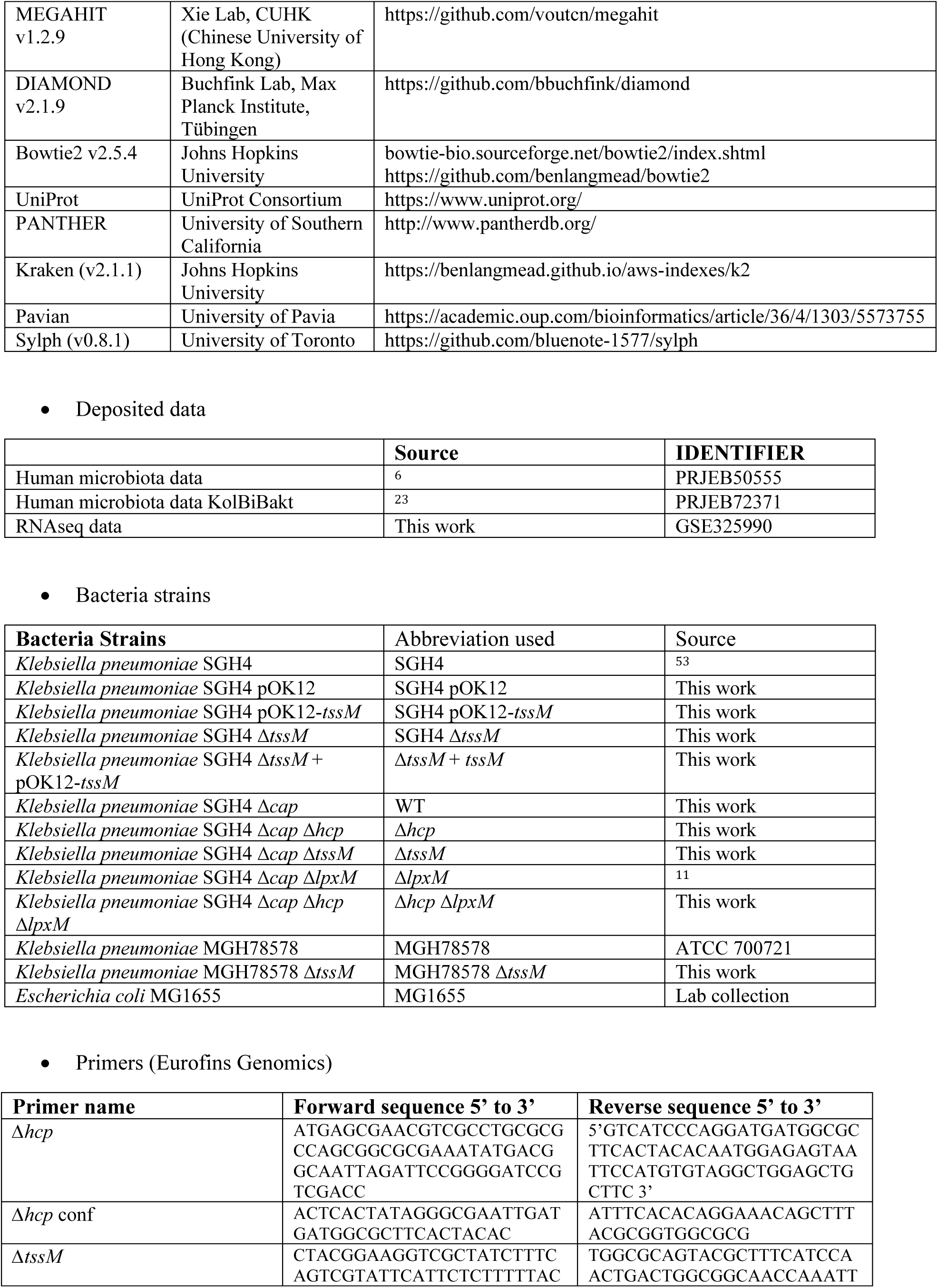
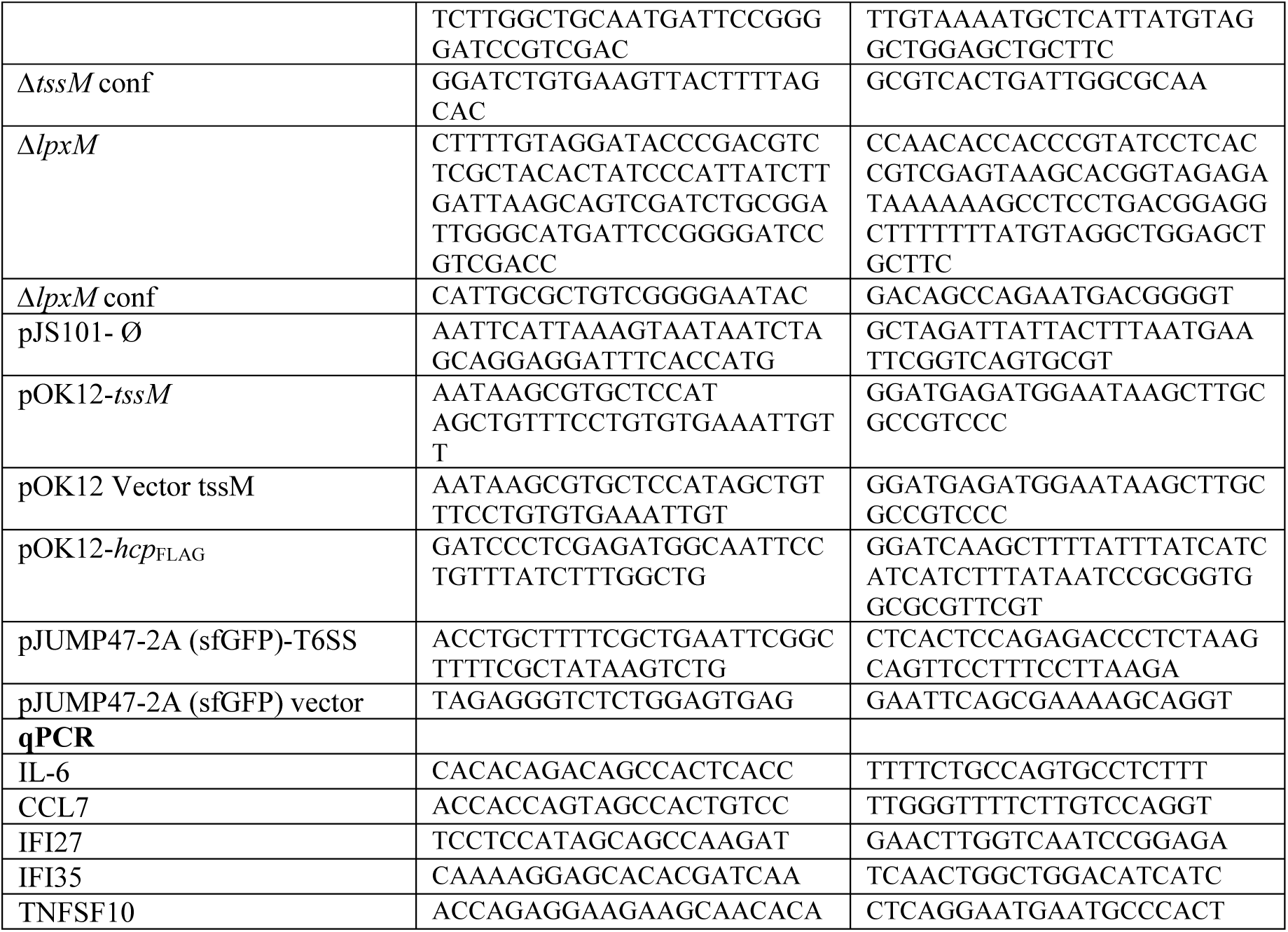

### Experimental model and subject details

#### Metagenomic Data Acquisition

Publicly available shotgun metagenomic sequencing datasets from two independent inflammatory bowel disease (IBD) cohorts were downloaded and analyzed: the Swedish KolBiBakt cohort (accession number PRJEB72371) and the French and Israeli cohorts (accession number PRJEB50555) described by Federici et al^6,23^. Only raw sequencing reads were used in the analyses and only samples with available clinical metadata (diagnosis and disease activity state: healthy control, remission, or flare) and CRP levels were included.

#### Mouse strains

All animal experiments were approved by the local ethical committee (Stockholms Norra djurförsöksetiska nämnd).

C57BL/6 mice were purchased from Jackson Laboratory and housed under specific pathogen-free (SPF) conditions at the Karolinska Institutet. Six-to eight-week-old adult females were used for the experiments. To reduce baseline microbiota, all mice were treated with 0.5 g/L streptomycin and 0.1 g/L amoxicillin in drinking water for three to five days prior to *K. pneumoniae* gavage.

#### Zebrafish strains

Transgenic zebrafish (*Danio rerio*) *Tg*(mpeg1:gal4-UAS:NTR-mCherry)^50^ expressing mCherry in macrophages were bred at the Karolinska Institute core facility. Fertilized embryos were raised in Petri dishes containing E3 medium (5 mM NaCl, 0.17 mM KCl, 0.33 mM CaCl_2_, 0.3 mM MgSO4) and immersed in 30 mg/l 1-phenyl-2-thiourea (PTU) in E3 from approximately 2-6 hpf for the duration of the experiment.

#### Bacterial strains

All bacterial strains used in this study are listed in the Key Resource Table. Bacteria were grown at 37°C under aerobic conditions on blood agar plates or in liquid media as indicated: DMEM supplemented with 10% FBS (Gibco), Luria Broth (LB) or low-salt LB. When stated, the following antibiotics were added: kanamycin (50 μg/mL), hygromycin (50 μg/mL), apramycin (50 μg/mL), and spectinomycin (100 μg/mL).

*K. pneumoniae* Δ*hcp*, Δ*tssM* and Δ*lpxM*Δ*hcp* mutants were generated using the λ red system as previously described^11^. Briefly, the gene of interest was replaced by an apramycin resistance cassette from pIJ773 in *K. pneumoniae* carrying the recombination system on pSIM18-hyg. The cassette was then removed using pFLP-hyg. The pOK12-*tssM* and pOK12-*hcp^FLAG^*plasmids were constructed by insertion of the XbaI-HindIII PCR fragment into pOK12 digested by the same enzymes. The pJS101-empty was obtained by excising the *GFP* gene using overlapping PCR. For transcriptional reporter construction, a 500 pb fragment encompassing the T6SS promoter was cloned in pJUMP47-sfGFP using overlapping PCR to create the pJUMP47-sfGFP-T6SS. Primers used for construction are listed in the Key Resource Table.

#### Human cell lines

BJ-hTERT human immortalized fibroblasts were cultured in DMEM supplemented with 10% FBS. THP-1 human monocytes (ATCC; TIB-202) were grown in RPMI (Gibco) supplemented with 10% FBS. THP-1 were differentiated into macrophage-like cells by 20 ng/mL phorbol 12-myristate 13-acetate (PMA; Sigma-Aldrich) treatment for 24 h, followed by 24 h rest in PMA-free medium prior to experiments. BJ-hTERT and THP-1 NF-κB reporter cells were generated using NF-κB Firefly Luciferase Reporter Lentivirus (Kerafast; FCT176) and selected 48 h with 2 μg/mL puromycin. Mycoplasma contamination was tested monthly using MycoAlert Mycoplasma Detection Kit (Lonza) according to the manufacturer’s instructions. All experiments were performed within 10 passages from frozen stocks.

## Method details

### T6SS signature and metagenomic analysis

The most conserved core Type VI Secretion System (T6SS) genes *tssA, tssB, tssK* and *tssL* from Enterobacteriaceae species and*, tssB*, *tssK* from Bacteroides were translated into protein sequences to generate a reference database. The resulting protein database was used for downstream homology searches.

All metagenomes were individually assembled using MEGAHIT v1.2.9 with default parameters. Contigs from each assembly were used for T6SS gene identification. Assembled contigs were queried against the T6SS protein database using DIAMOND v2.1.9 (blastx, sensitive mode). Contigs matching *tssA, tssB, tssK* or *tssL* were extracted into separate FASTA files for each metagenome. Subsequence coordinates corresponding to each T6SS gene were retrieved from DIAMOND output TSV files, and the corresponding contig regions were extracted. Raw metagenomic reads were mapped back to the extracted T6SS sequences using Bowtie2 v2.5.4 with default settings. Mapped read pairs were output as FASTQ files. Read counts mapping to each T6SS gene were then quantified to estimate gene abundance. Normalization for sequencing depth was applied prior to downstream analyses.

The taxonomy of the T6SS mapping reads was assigned using Kraken (v2.1.1) with the standard database (version of June 2024, capped at 8Gb). Family-level taxonomic abundance tables were generated using Pavian imported into R (v4.x) and converted to relative abundances by normalizing each sample to 100%. Stacked bar plots showing relative abundance per condition were generated using the *ggplot2* package.

The full taxonomic profiling of all metagenomics reads was performed using Sylph (v0.8.1) with the GTDB (r226). Relative abundance of *K pneumoniae* was obtained from taxonomic profiling of shotgun metagenomes. T6SS abundance was quantified as normalized gene counts (reads per million, RPM). Associations between *Klebsiella* relative abundance and T6SS abundance were assessed using Spearman’s rank correlation to accommodate non-normal, zero-inflated distributions. Correlations were computed across all samples and stratified by disease type (Crohn’s disease or ulcerative colitis) and clinical activity (flare or remission). Beta diversity analyses, including Bray-Curtis principal coordinates analysis (PCoA), were performed based on the taxonomic profiles generated by Sylph. Statistical tests were performed using R (v4.x). P values < 0.05 were considered significant.

### Zebrafish microinjection and imaging

Zebrafish larvae were anesthetized with 0.016% tricaine and positioned on an agarose injection mold. A 1 nL inoculum containing 1,000 CFU of *K. pneumoniae* carrying the pJS102 plasmid (constitutively expressing GFP), 0.05% phenol red suspended in PBS was microinjected into the otic vesicle using a pulled glass capillary needle and a microinjector. Control larvae received an equivalent volume of 0.05% phenol red in PBS. Following injection, larvae were screened for correct injection with a fluorescence microscope, and individual larvae were transferred to 96-well plate containing exposure medium (E3, PTU and Tricaine) and imaged using the ACQUIFER Imaging Machine (Bruker). CFU consistency within and between experiments was followed with frequent platings on LB agar plates (data not shown). Z-stack images were acquired every 20 minutes for 17 hours with a 4X objective in GFP, mCherry, and brightfield channels. Imaging parameters (exposure, gain, laser intensity) were kept constant across all samples. Image stacks were processed in Fiji/ImageJ (National Institute of Health, USA). The otic vesicle was manually outlined, and the same region of interest (ROI) size was applied across all timepoints image stacks. *K. pneumoniae* growth and clearance as well as macrophages recruitment to the otic vesicle were assessed by measuring GFP and mCherry fluorescence intensity using the *Measure* function, recording integrated density and mean fluorescence intensity. Background fluorescence was subtracted from each image before analysis. Statistical significance was assessed using one-way ANOVA followed by Tukey’s multiple comparisons test.

### DSS-induced colitis model

Mice were orally inoculated with 10^9^ CFU of either WT or Δ*hcp K. pneumoniae* strains carrying the pJS101-Ø plasmid one day after antibiotic treatment. 24 h post-gavage, colitis was induced by administrating 2% DSS in drinking water ad libitum for 7 days followed by a one-day recovery period prior to sacrifice. Mice were monitored daily for weight loss and health status over the 15-day experiment. Fecal samples were collected daily, homogenized, and plated on spectinomycin LB plates to quantify *K. pneumoniae* colonization.

### AOM/DSS model of colitis-associated cancer

AOM (10 mg/kg) was administered to mice by intraperitoneal injection prior to 3 days of antibiotic treatment, then mice were subjected to three rounds of DSS. For each DSS cycle, mice were orally inoculated with 10^9^ CFU of either WT or Δ*hcp K. pneumoniae* strains carrying the pJS101-Ø plasmid one day prior to DSS administration and again on day 3 of DSS treatment. Mice received 2% DSS in drinking water *ad libitum* for five days, followed by a 14-days recovery period. Throughout the experiment, mice were weighed and monitored daily to assess disease progression. Fecal samples were collected before and after each bacterial gavage, and weekly during the recovery phases. At the experimental endpoint, colon tissues were harvested for further analysis. Macroscopically visible tumors were counted, and tumor volume was estimated assuming a spherical shape.

### Histological preparation and scoring

Mouse colons were Swiss-rolled, and fixed in 10% neutral-buffered formalin, and processed for paraffin embedding. Sections (4 μm thick) were stained with hematoxylin and eosin (H&E) and inflammation was scored in a double-blind manner based on histopathological features as previously described^54^.

For immunohistochemistry (IHC), tissue sections were subjected to citrate-based antigen retrieval followed by incubation with anti-Ki67 or anti-p65 antibody (1:100 dilution), followed by a biotinylated secondary antibody (1:1000 dilution). Detection was performed using the ABC-peroxidase method with 3,3’-diaminobenzidine (DAB) as a chromogen. Stained sections were imaged using an AxioScan.Z1 Slide Scanner (Zeiss). The percentage of Ki-67-or p65-positive nuclei relative to total nuclei was calculated for each sample.

### Multiplex immunofluorescence

mIF was performed at the Spatial Proteomics National Facility (SciLifeLab, Stockholm, Sweden) on a COMET platform (Lunaphore Technologies), as previously described^55^. Antibodies are listed in the Key Resource Table. Blank images of the TRITC and Cy5 channels were captured prior to staining for autofluorescence subtraction. A 1-minute blocking step with BU10 (2.5% normal goat serum) preceded each cycle. Primary antibodies (in 0.05% TBS-T) were incubated for 8 minutes, and secondary antibodies (TRITC: Alexa Fluor goat anti-rat AF555, 1:100; Cy5: Alexa Fluor 647 goat anti-rabbit, 1:200 in 0.05% TBS-T) for 8 minutes, except cycle 7 (F4/80), where incubation was 4 minutes. DAPI was co-stained in every cycle for image alignment. Images were acquired using a CMOS IDS UI-3280CP-M-GL camera (2456 × 2054 px; 0.23 μm per pixel) with a 20X objective (0.75 NA/1 mm WD). Exposure times were optimized for TRITC and Cy5; DAPI exposure was 25 ms. Antibodies were eluted with BU07 for 4 minutes per cycle (2 minutes for cycle 7). This process was repeated for all antibody cycles, CD3 and FOXP3 staining was performed manually following the 8-plex protocol. All samples were prepared and stained in batches of four.

### Spatial profiling of immune cells

mIF images were analyzed using QuPath^56^ to quantify immune cell populations. Image alignment was performed using DAPI as a reference channel. Automated cell detection was applied to segment nuclei and assign cell boundaries based on fluorescence intensity, as previously described^55^. A machine learning classifier was trained in QuPath using morphological features to distinguish normal tissue, low-grade dysplasia, and high-grade dysplasia. Trained classifiers for immune cell populations and tissue features were integrated to enable automated annotation of immune cells across tissue regions and samples. Quantification of total and co-stained cell populations was then performed to assess immune marker distribution and spatial organization across normal and dysplastic tissues.

### ELISA and cytokine array

Fecal samples from individual mice were homogenized in PBS containing protease inhibitors and centrifuged at 12,000 × g for 10 min. Fecal Lipocalin-2 (Lcn-2) levels were determined by ELISA (Invitrogen, EMLCN2). Blood collected at endpoint was centrifuged at 10,000 × g for 10 min at 4°C to isolate serum, and IL-6, IFN-α and IFN-γ were quantified by ELISA (Invitrogen KMC006; BMS6027; KMC4021), following the manufacturer’s instructions. Systemic cytokine profiles were assessed using a mouse cytokine antibody array for 96 cytokines (ab193659, Abcam). Briefly, pooled serum samples were diluted, incubated with the array membrane overnight at 4°C, washed, and then probed with a biotin-conjugated antibody cocktail followed by HRP-conjugated streptavidin. Chemiluminescent signals were detected using a ChemiDoc imaging system (Bio-Rad).

### Cell infections and treatments

For transient infections, overnight cultures of *K. pneumoniae* were grown in DMEM or RPMI supplemented with 10% FBS, then diluted in fresh medium. BJ-hTERT or macrophage-like THP-1 cells were infected at a multiplicity of infection (MOI) ranging from 10 to 100, as indicated, for 30 min (Western blots) or 4 h (RT-qPCR). Transwell inserts (0.4 µm pore size, Corning) were used to assess the effects of secreted bacterial factors while preventing direct bacterial-host cell contact.

Bacterial cell-free supernatants were prepared from *K. pneumoniae* cultured in DMEM without phenol red or in RPMI at 37°C with shaking (200 rpm) until optical density (OD_600_) reached 0.8. Bacteria were centrifuged at 4,000 rpm for 10 min, and the supernatant was filtered through a 0.22 µm pore-size filter (Filtropur S; Sarstedt) and stored at-20°C until use. For treatments, *K. pneumoniae* supernatant was added to cells at 1:100 dilution for 30 min (Western blot) or 4 h (RT-qPCR). For TLR4 inhibition, cells were pre-treated 2 h with 1μM TAK-242 (Sigma-Aldrich) before addition of bacterial supernatant. After treatment, cells were washed three times with PBS and stored at-80°C until further analysis.

### RNA sequencing

Macrophage-like THP-1 cells were infected with *K. pneumoniae* WT or Δ*tssM* strains for 4 h in three independent biological replicates. Total RNA was extracted using Aurum total RNA kit (Bio-Rad). RNA-seq was performed by Vertis Biotechnologie AG. Random-primed cDNA libraries were prepared following Illumina protocol. The cDNA pool was paired-end sequenced on an Illumina HighSeq system using 2×150 bp read length. Quality control was performed using FastQC. Reads were processed with the nf-core/rnaseq pipeline (v3.1)^57^. HISAT2 (v2.1.0) was used to align the raw RNA-seq fastq reads to the human reference genome (GRCh38.97). Read quantification was computed using featureCounts (v1.6.4). Differential analysis was performed with the R package DESeq2. False positive discovery (FDR)<0.05 was set as cut-off for DEGs. Gene ontology (GO: biological process) and pathway enrichment analysis were performed using g:Profiler^52^ and visualized using Cytoscape (v3.6.1)^58^ as described by Reimand *et al*.^59^. Enrichment analysis of transcription factors binding motifs (from TRANSFAC database) in DEGs promoters was performed using g:Profiler. GSEA was performed using GSEA software (https://www.gsea-msigdb.org/gsea/index.jsp)^60^. The complete lists of DEGs and enriched GO terms are provided in Supplementary Table S1.

### RT-qPCR

RNA was extracted using Aurum total RNA kit (Bio-Rad), and cDNA was synthesized using iScript cDNA synthesis kits (Bio-Rad). qPCR was performed with SsoAdvanced Universal SYBRGreen SuperMix (Bio-Rad) on a CFX384 Real-Time System (Bio-Rad). qPCR primers are detailed in the Key Resource Table. *RPL13A* and *GAPDH* were used as reference genes. Data analysis was processed using Bio-Rad CFX 3.1 using the ΔΔCT method. Error bars represent standard deviation from mean of at least three independent experiments.

### Protein extraction and western blotting

For human cell lysates, proteins were extracted using ice-cold RIPA buffer (Thermo Fisher) supplemented with cOmplete protease and PhosSTOP phosphatase inhibitors (Roche). Protein concentration was determined using the BCA Protein Assay Kit (Thermo Fisher). Bacterial proteins were extracted by bead beating (Matrix B, MP Biomedicals) for two cycles of 30 sec in RIPA supplemented with cOmplete protease inhibitors, then mixed with Laemmli buffer to a final concentration of 0,5 OD₆₀₀ per 10 µl. For the extracellular fraction, proteins from bacteria-free supernatant were precipitated in 10% (v/v) trichloroacetic acid (TCA) overnight at 4°C followed by 1 h centrifugation at 15,000 × g, 4°C. Protein pellets were washed with ice-cold acetone and air-dried, then resuspended with RIPA-Laemmli buffer to a final concentration of 2 OD₆₀₀ per 10 µl. For Western blots, samples were denatured in Laemmli buffer at 95°C for 5 min and separated by SDS-PAGE. Western blot signals were detected by chemiluminescence using a ChemiDoc Imaging System (Bio-Rad). Antibodies are listed in the Key Resource Table.

### NF-κB luciferase reporter assay

BJ-hTERT and differentiated THP-1 NF-κB reporter cells seeded in 96-well plate were either infected with *K. pneumoniae* at MOI 10 or treated with bacterial supernatant (1:100) for 5 h. Luciferase activity was measured using Firefly Luciferase Glow Assay Kit (Pierce) according to supplier’s instructions.

### T6SS operon transcription reporter assay

*K. pneumoniae* strains carrying the T6SS promoter-*sfgfp* fusions (pJUMP47-sfGFP-T6SS) were cultured in either LB or phenol red-free DMEM to an OD_600_ of 0.8. GFP fluorescence was measured using a SparkCyto plate reader (Tecan). Background fluorescence from bacteria-free medium was subtracted, and fluorescence intensity (RFU) was normalized to OD_600_ (CFU) to account for differences in bacterial density.

### Bacterial competition assays

Bacterial competition assays were performed similarly as previously described^27^. Briefly, attacker and prey strains were grown overnight in LB or DMEM at 37°C with shaking, washed, and adjusted to an OD_600_ of 0.5. Bacteria were mixed at a 1:1 ratio and spotted onto pre-warmed blood or LB agar plates for 3-6 hours at 37°C, then recovered in PBS, serially diluted, and plated on selective media to quantify prey survival. T6SS-deficient mutants were used to confirm T6SS-dependent killing.

### LPS, lipid quantification and LPS structure analysis

For LPS quantification, cell-free supernatants were diluted (1:5,000 to 1:25,000) in pyrogen-free water, and LPS levels were measured using the Limulus Amebocyte Lysate (LAL) Chromogenic Endotoxin Quantification Kit (Pierce) following manufacturer’s instructions. Enzymatic units of reactogenic LPS were normalized to the OD₆₀₀ of the bacterial culture. For lipid quantification, 25 μL of *K. pneumoniae* culture supernatant (OD_600_ = 0.8) was mixed with 200 μL of FM4-64 dye (2.25 μg/mL) and incubated for 10 min at 37°C. Fluorescence was measured using a SparkCyto plate reader (Tecan) (Ex: 485 nm / Em: 670 nm). Background fluorescence from fresh medium was subtracted. All data are presented as mean ± SEM from at least three independent biological replicates. For LPS structure analysis, LPS was extracted from *K. pneumoniae* strains using LPS extraction kit (Sigma-Aldrich; MAK339) according to the manufacturer’s instructions. LPS loading was normalized to protein concentration (800ng) and loaded on NuPAGE™ Bis-Tris Mini Protein Gels, 4–12%, 1.0mm and stained with Imperial Protein stain.

### OMVs isolation and analysis

Cell-free supernatants from *K. pneumoniae* cultures in phenol red-free DMEM (OD_600_ = 0.8) were enriched in OMVs by ultrafiltration through Vivaspin columns (100 kDa). OMVs size distribution was determined on the enriched fraction by DLS on a Zetasizer Ultra instrument (Malvern). OMVs were then isolated as previously described^61^ using OptiPrep density gradient ultracentrifugation at 155,000 × g, 4°C for 8 h and resuspended in PBS. OMVs protein profiling was performed by SDS-PAGE followed by silver saining using the Pierce Silver Stain Kit, according to the manufacturer’s instructions. For protein identification, OMVs were analyzed by LC-Orbitrap MS/MS at the Mass Spectrometry Based Proteomics Facility, Uppsala University, using a 90-minute gradient acquired mass spectra were analyzed using Proteome Discoverer 1.4 and searched against the *K. pneumoniae* SGH10 proteome from UniProt (https://www.uniprot.org/). Identified proteins subcellular localizations were assigned using UniProt and PANTHER (http://www.pantherdb.org/)^62^.

## Statistical Analysis

Statistical analyses were performed using GraphPad Prism. Unless otherwise stated, statistical significance was assessed two-using tailed unpaired t test on at least three independent biological replicates. * *p* < 0.05; ** *p* < 0.01; *** p < 0.001; **** p < 0.0001.

---

**Supplementary Figure S1.**
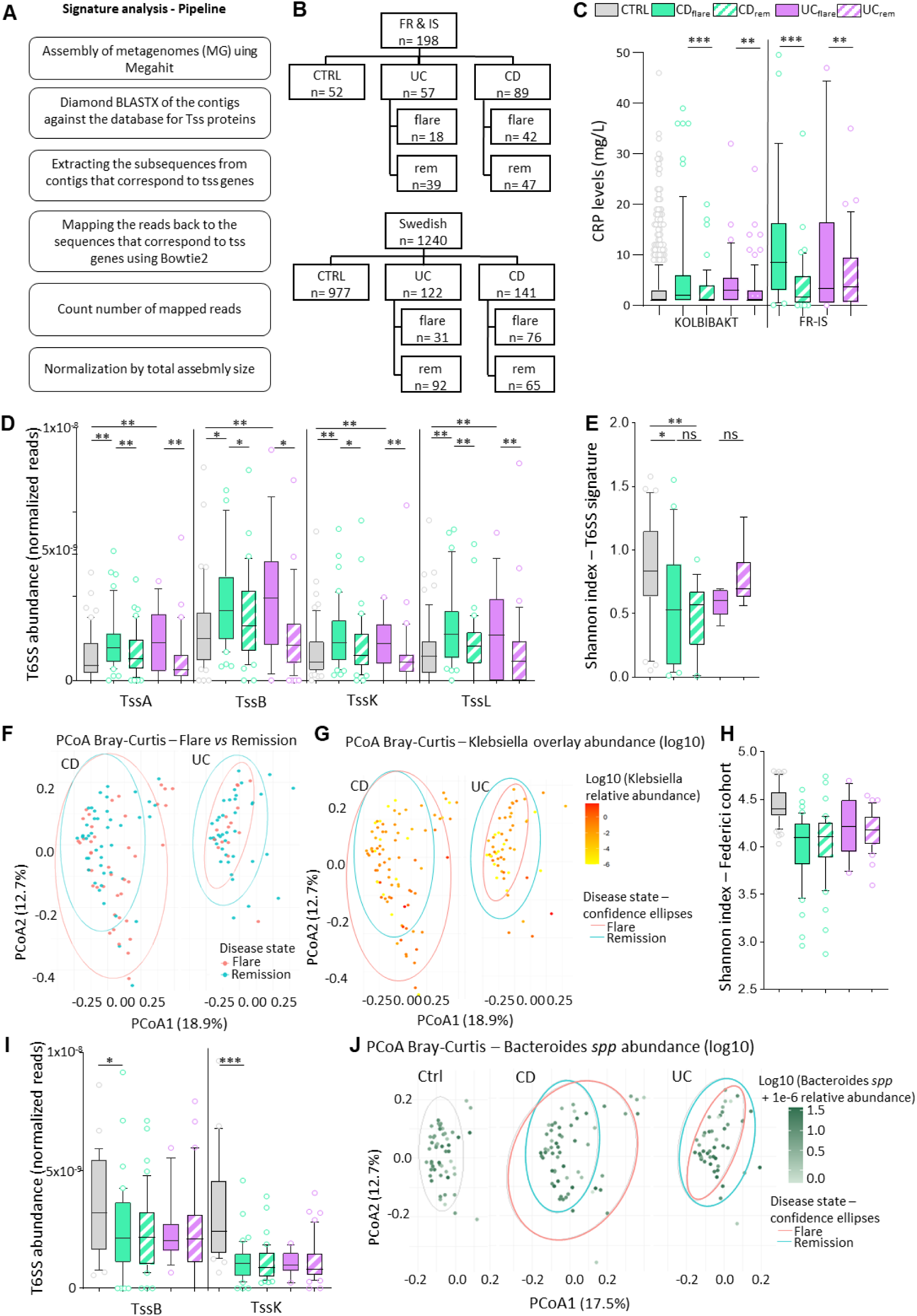
Overview of metagenomic pipeline, cohort composition and microbial analyses. (A) Schematic overview of the metagenomic signature analysis pipeline. (B) Flowchart of sample distribution across cohorts, indicating numbers of controls (CTRL), ulcerative colitis (UC), Crohn’s disease (CD). (C) Comparison of CRP levels (mg/L) across CTRL, CD_high_, CD_low_, UC_high_, and UC_low_ groups. Box plots show median, and individual samples. p-values: *p < 0.05, **p < 0.01, ***p < 0.001. (D) Abundance of T6SS structural genes (*tssA*, *tssB*, *tssK*, *tssL*) normalized to total mapped reads across cohorts. Box plots show median and individual data points. Statistical comparisons as in panel C. (E) Shannon diversity index across CTRL and IBD subgroups performed on the families identified in the T6SS signature. ns, not significant. (F-H) Cohort community structure analyses. (F-G) Bray–Curtis dissimilarity analysis of microbial community profiles across groups (F), showing *Klebsiella* overlay abundance in log10 (G). (H) Shannon diversity index calculated across the full species set of the entire cohort. (I-K) T6SS gene abundance and *Bacteroides*-focused analyses. (I) Normalized abundance of T6SS genes (*tssB*, *tssK*) across cohorts. Statistical annotations as in panel C. (J) Relative abundance of *Bacteroides* spp. across the cohort.

**Supplementary Figure S2.**
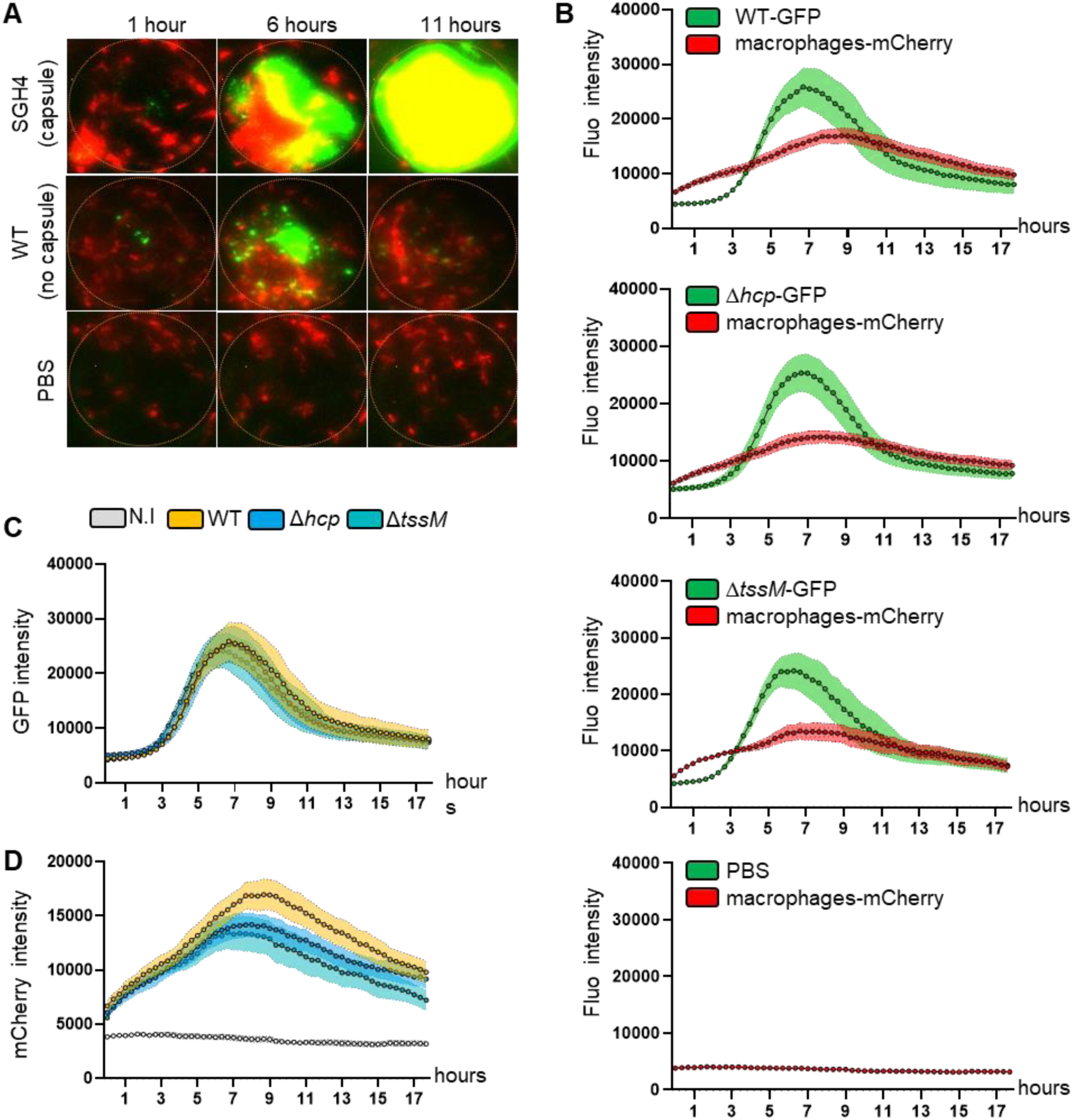
Time-lapse imaging of macrophage recruitment and *K. pneumoniae* dynamics in zebrafish otic vesicles, related to. **Figure 2**. (A) Representative time-lapse images of zebrafish otic vesicles microinjected with GFP-labeled *K. pneumoniae* SGH4 strains, either capsulated (SGH4 capsule) or non-capsulated (WT), showing bacterial dynamics and mCherry-labeled macrophage recruitment over time. (B) Quantification of fluorescence intensity for GFP-labeled bacteria and mCherry-labeled macrophages over an 18-hour period (images acquired every 20 min). (C) Quantification of bacterial burden (GFP fluorescence intensity) in otic vesicles of non-infected controls (N.I.), WT-, and T6SS mutant–infected zebrafish larvae (Δ*hcp* and Δ*tssM*). (D) Quantification of macrophage recruitment (mCherry fluorescence intensity) under the same conditions.

**Supplementary Figure S3.**
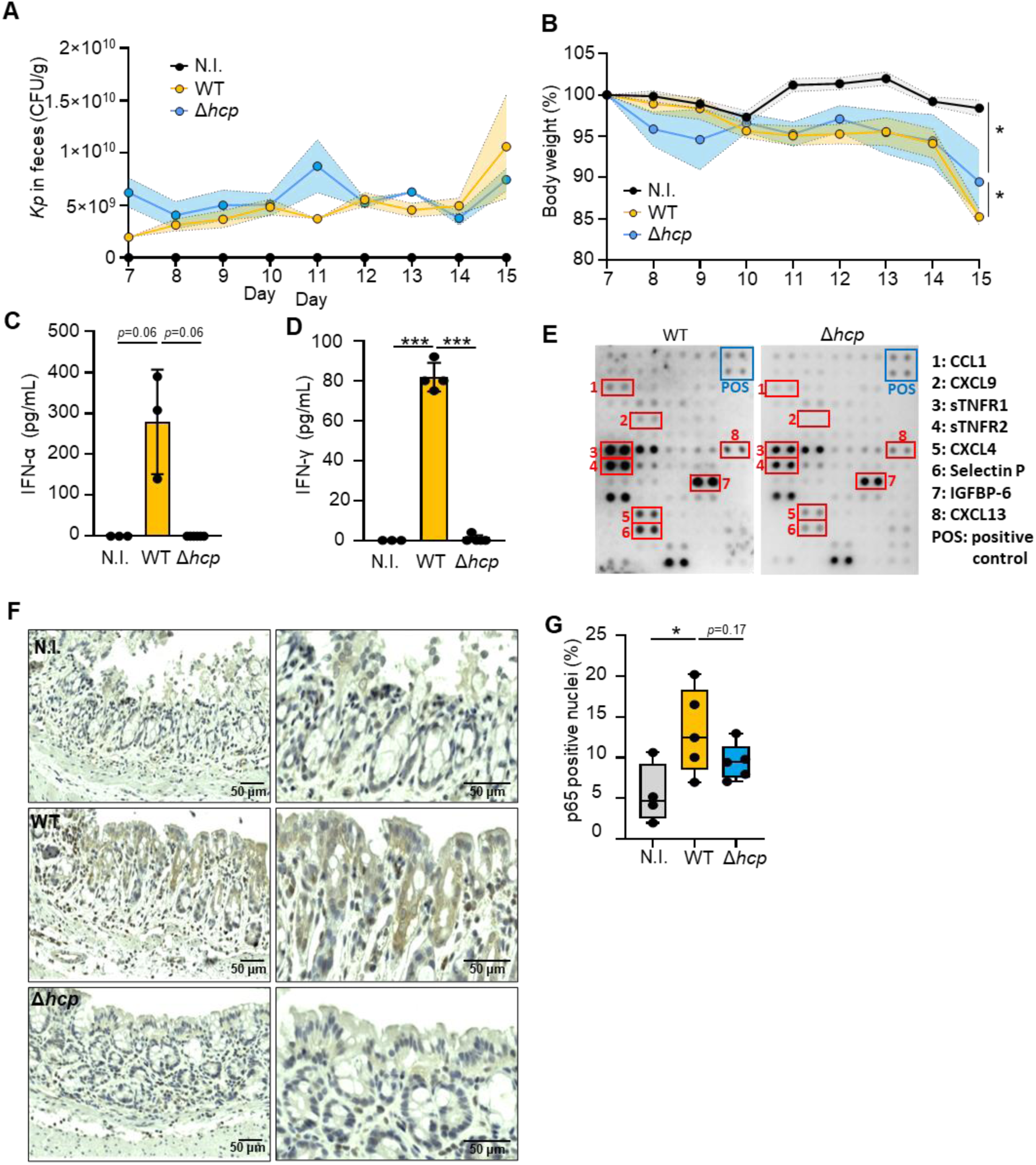
*K. pneumoniae* T6SS exacerbates inflammation without affecting bacterial load during mono-colonization in mice, related to. **Figure 2**. (A) Fecal *K. pneumoniae* colonization levels (CFU/g) over time in DSS-treated mice colonized with WT or Δ*hcp* strains. N.I.: not infected. (B) Mice body weight during the course of DSS treatment and colonization. * p < 0.05 (C–D) Serum levels of type I and type II interferons measured by ELISA: IFN-α (C) and IFN-γ (D). Error bars ± SD. *** p < 0.01. (E) Cytokine array of serum collected at sacrifice comparing WT-and Δ*hcp*-colonized mice. (F) Representative immunohistochemistry (IHC) images of p65 staining in colon Swiss roll sections. (G) Quantification of p65-positive nuclei in colonic cells. Error bars ± SD. * p < 0.05.

**Supplementary Figure S4.**
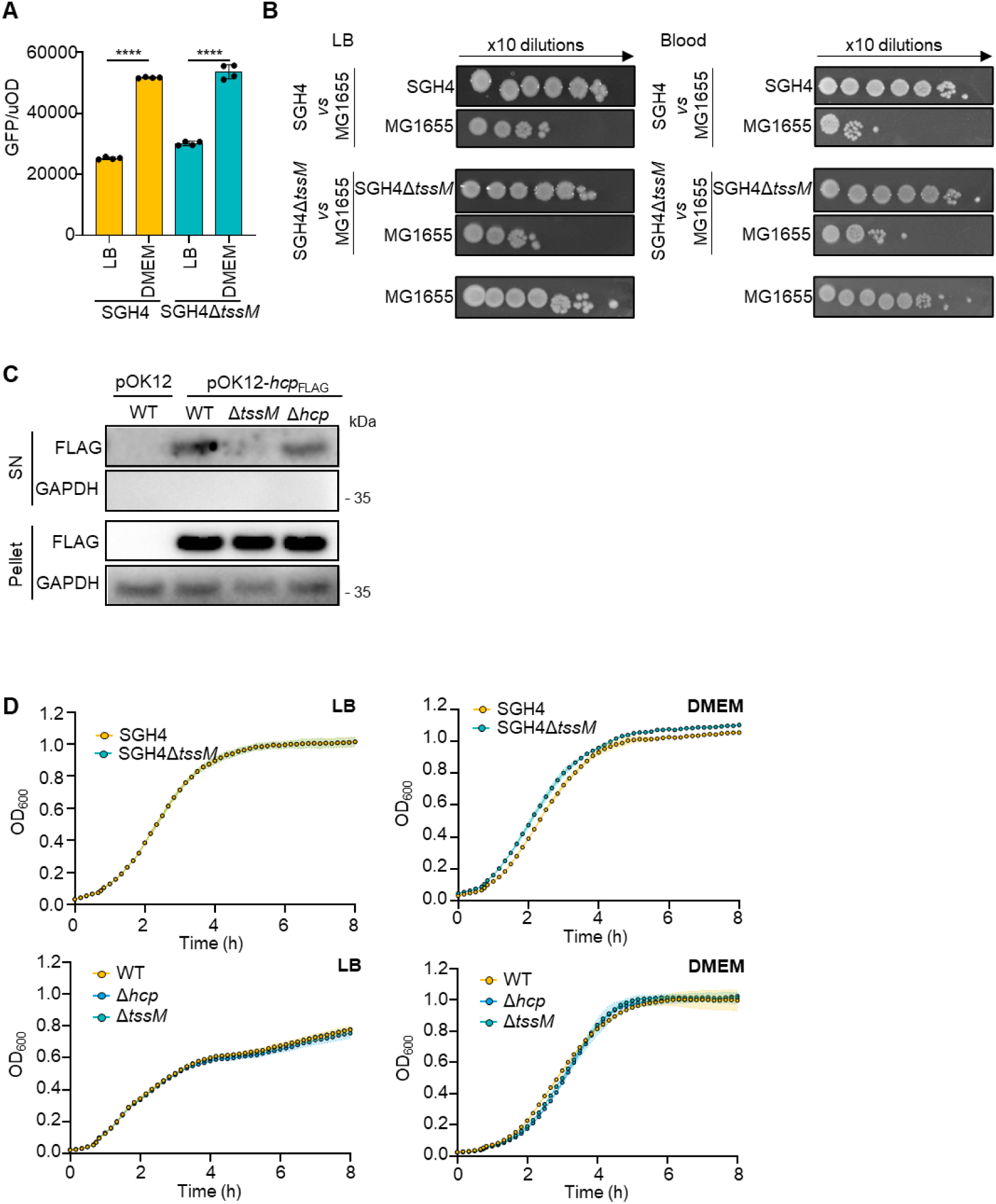
Functional validation of *K. pneumoniae* T6SS, related to. **Figure 3**. (A) T6SS promoter activity in SGH4 and SGH4 Δ*tssM* strains carrying the pJUMP43-sfGFP reporter plasmid, in which GFP is expressed under control of T6SS promoter. GFP fluorescence was normalized to OD_600_ and measured in both LB and DMEM media. Error bars ± SD. **** p < 0.0001. (B) T6SS-dependent antibacterial activity assessed by bacterial competition assays on LB and blood agar plates, using SGH4 or SGH4 Δ*tssM* as attacker strains and *E. coli* MG1655 as prey. (C) Hcp secretion assay in strains harboring pOK12-*hcp*^FLAG^. FLAG-tagged Hcp was detected by western blot in both supernatant (SN) and pellet fractions, with GAPDH used as a control for cell lysis and loading. (D) Growth curves of SGH4, SGH4 Δ*tssM*, WT, Δ*hcp*, and Δ*tssM* strains in LB and DMEM media showing comparable fitness under experimental conditions.

**Supplementary Figure S5.**
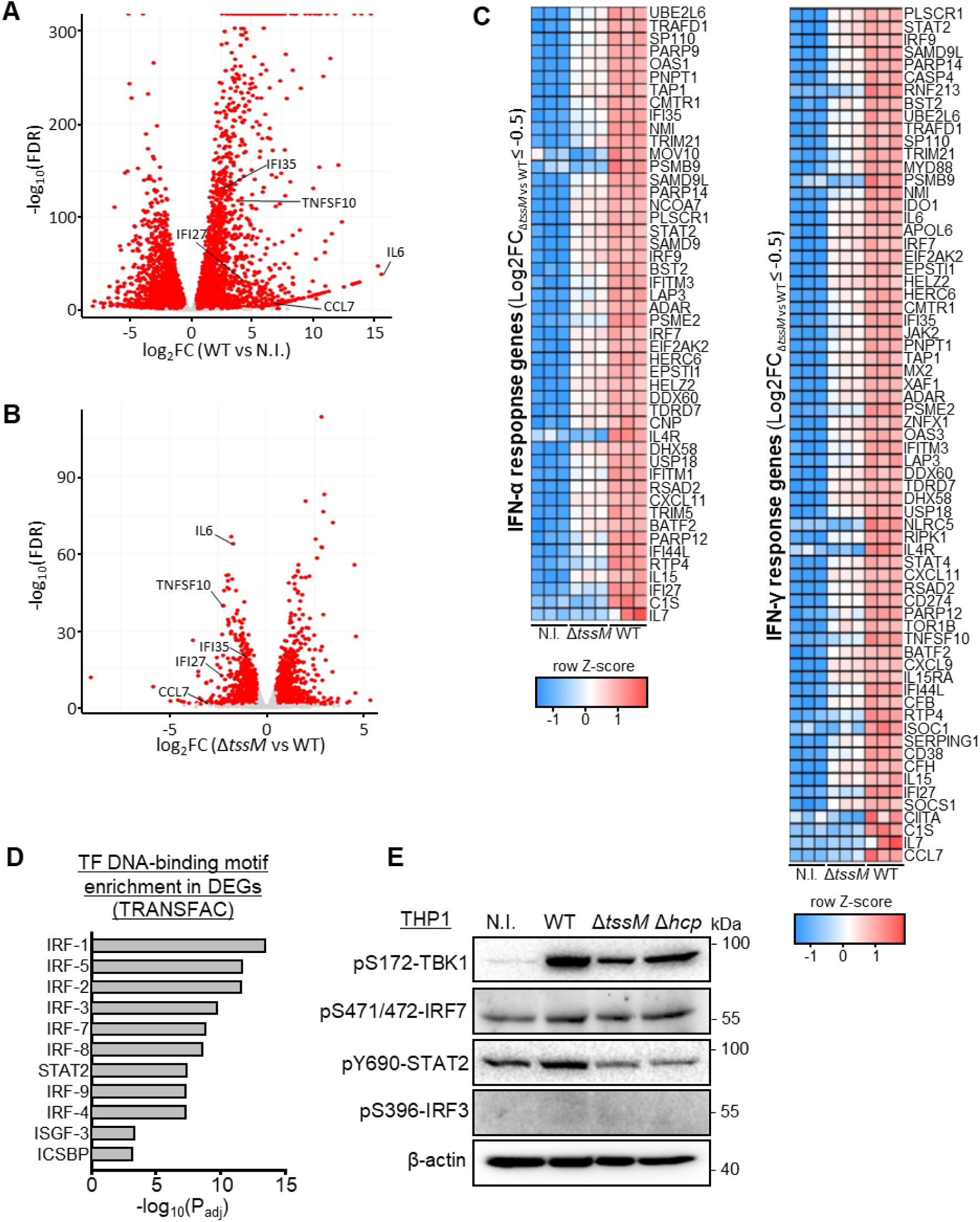
Transcriptomic analysis of THP-1 cells infected with *K. pneumoniae* WT and Δ*tssM* strains, related to. **Figure 3**. (A–B) RNA-seq analysis DEGs comparing WT-infected versus non-infected (N.I.) THP-1 cells (A); and Δ*tssM*-infected versus WT-infected THP-1 cells (B). DEGs validated by RT-qPCR are highlighted. (C) Expression heatmap of DEGs in the response to interferon-α (GO:0035455) and interferon-γ (GO:0034341) pathways across conditions. (D) TRANSFAC analysis of top transcription factor DNA binding motifs enriched in DEGs between Δ*tssM*-infected and WT-infected THP-1 cells. (E) Western blot analysis of TBK1, IRF3, IRF7 and STAT2 activation using phospho-specific antibodies in THP-1 cells infected with *K. pneumoniae* WT, Δ*hcp*, or Δ*tssM* strains.

**Supplementary Figure S6.**
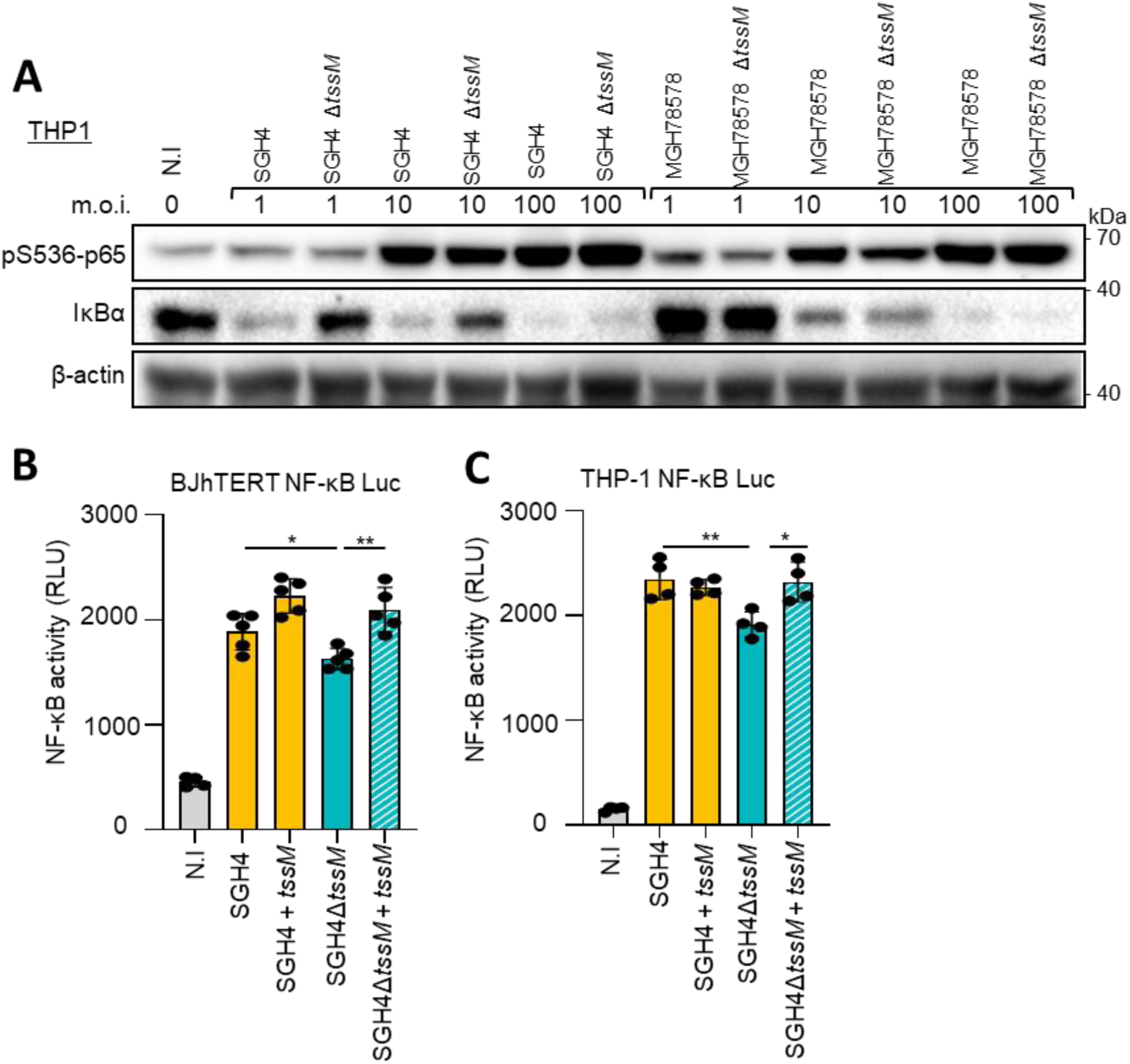
Complementation and strain validation confirm T6SS-dependent NF-κB activation, related to Figure 3. (A) Western blot analysis of NF-κB activation, showing p65 phosphorylation and IκBα degradation in THP-1 cells infected with *K. pneumoniae* SGH4 (capsule) and MGH7878 strains or their respective Δ*tssM* mutants across a range of MOIs (1 to 100). (B–C) NF-κB transcriptional activity in BJ hTert (A) and THP-1 (B) cells assessed by luciferase reporter assay. Cells were non-infected (N.I.) or infected with *K. pneumoniae* SGH4 or SGH4Δ*tssM* strains carrying either an empty plasmid (pBAD33-gent) or a complementation plasmid expressing *tssM* (pBAD33-gent-*tssM*). Complemented strains are denoted as SGH4 + *tssM* and SGH4*ΔtssM* + *tssM*. Error bars ± SD. * p < 0.05; ** p < 0.01.

**Supplementary Figure S7.**
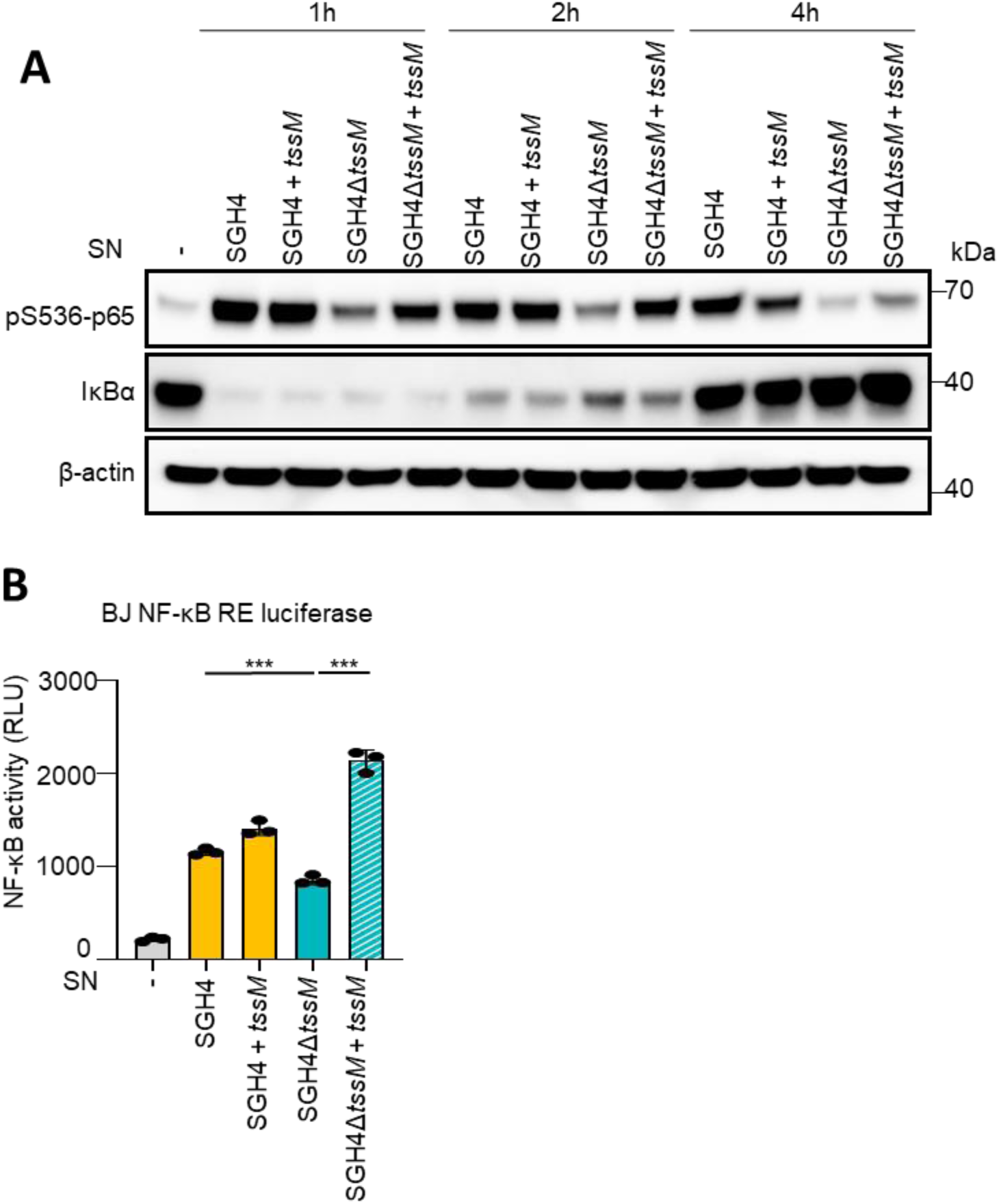
Complementation of T6SS restores NF-κB activation in response to bacterial supernatants, related to. **Figure 4**. (A) Western blot analysis of NF-κB activation, showing p65 phosphorylation and IκBα degradation in THP-1 cells treated for 1-4 h with bacterial supernatants (SN) from *K. pneumoniae* SGH4 (capsule) and SGH4Δ*tssM* strains carrying either an empty plasmid (pBAD33-gent) or a complementation plasmid expressing *tssM* (pBAD33-gent-*tssM*). Complemented strains are denoted as SGH4 + *tssM* and SGH4Δ*tssM* + *tssM*. (B) BJ hTert cells were treated with bacterial supernatants from SGH4 (capsule) or SGH4Δ*tssM* strains carrying either the empty vector or the complementation construct. NF-κB transcriptional activity measured by luciferase reporter assay. Error bars ± SD. *** p < 0.001.

**Supplementary Figure S8.**
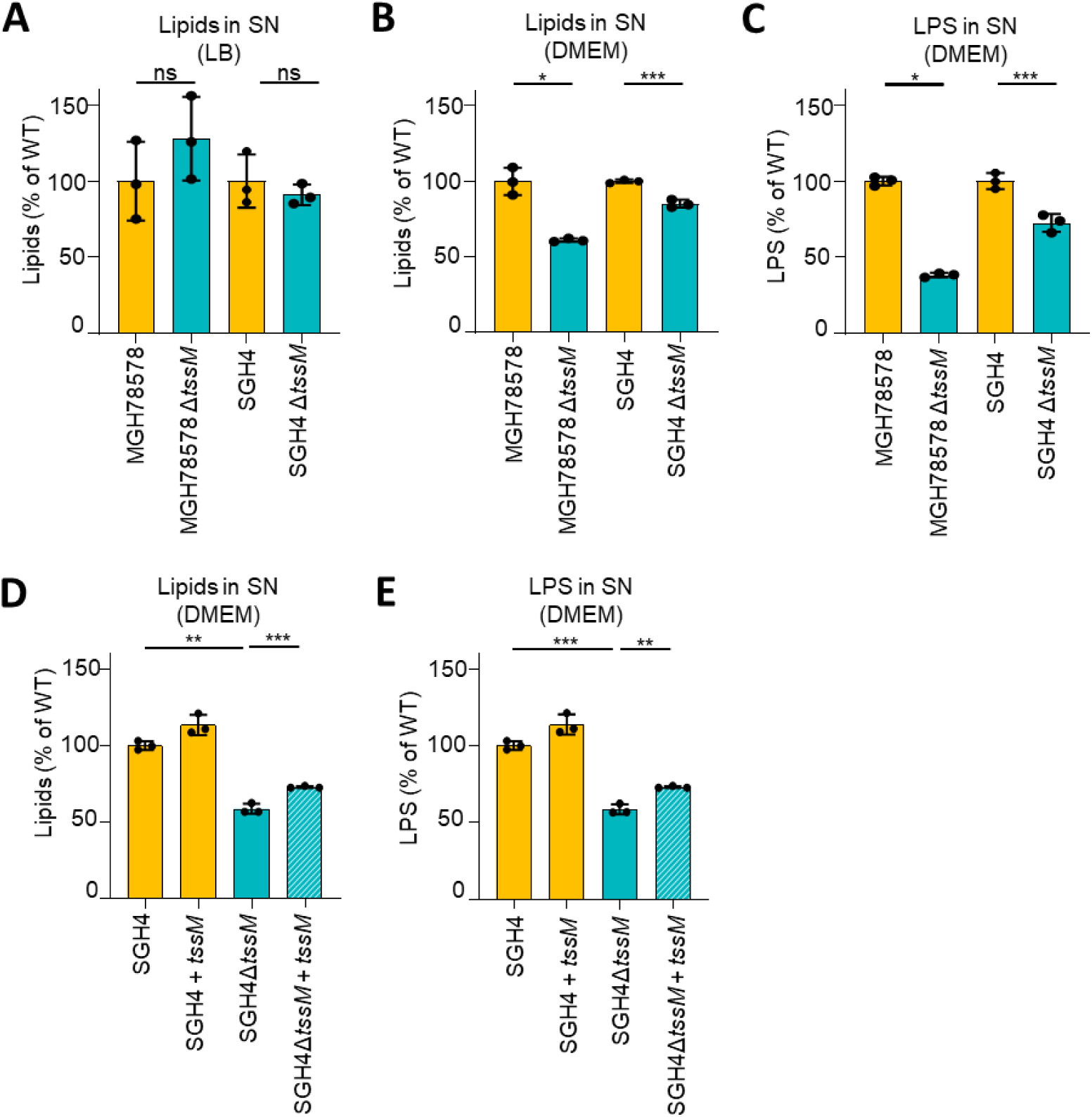
Complementation and strain validation confirm T6SS-dependent increase in lipid and LPS secretion, related to Figure 5. (A–B) Quantification of total lipid content using FM4-64 fluorescence in supernatants (SN) from *K. pneumoniae* SGH4 (capsule) and MGH78578 strains and their corresponding Δ*tssM* mutants cultured in LB (A) or DMEM (B). (C) LPS levels measured by LAL assay in supernatants from the same strains cultured in DMEM. (D–E) Complementation of T6SS activity in SGH4 Δ*tssM* using the *tssM*-expressing plasmid pBAD33-gent-tssM. Lipid quantification in SN by FM4-64 fluorescence (D). LPS secretion quantified by LAL assay in SN (E). Error bars ± SD. * p < 0.05; ** p < 0.01; *** p < 0.001; ns not significant.

**Supplementary Figure S9.**
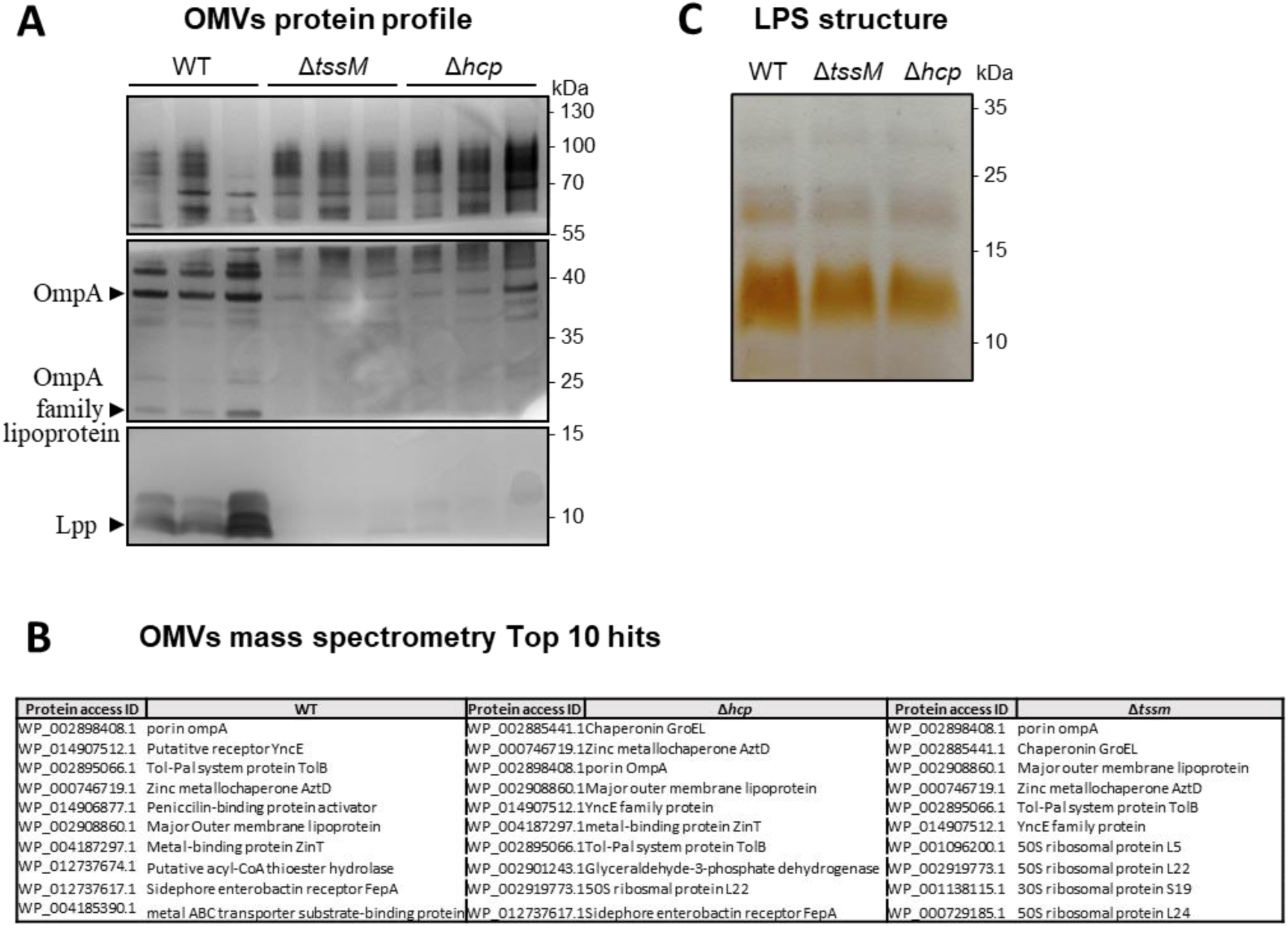
Analysis of OMVs and LPS composition in WT and T6SS mutant strains, related to Figure 5. (A) SDS-PAGE followed by silver staining of purified OMVs from WT, Δt*ssM*, and Δ*hcp* strains. Proteins indicated by arrows were excised and identified by mass spectrometry. (B) SDS-PAGE analysis of purified LPS from WT, Δ*tssM*, and Δ*hcp* strains. (C) Top protein hits identified by mass spectrometry for each strain.

**Supplementary Figure S10.**
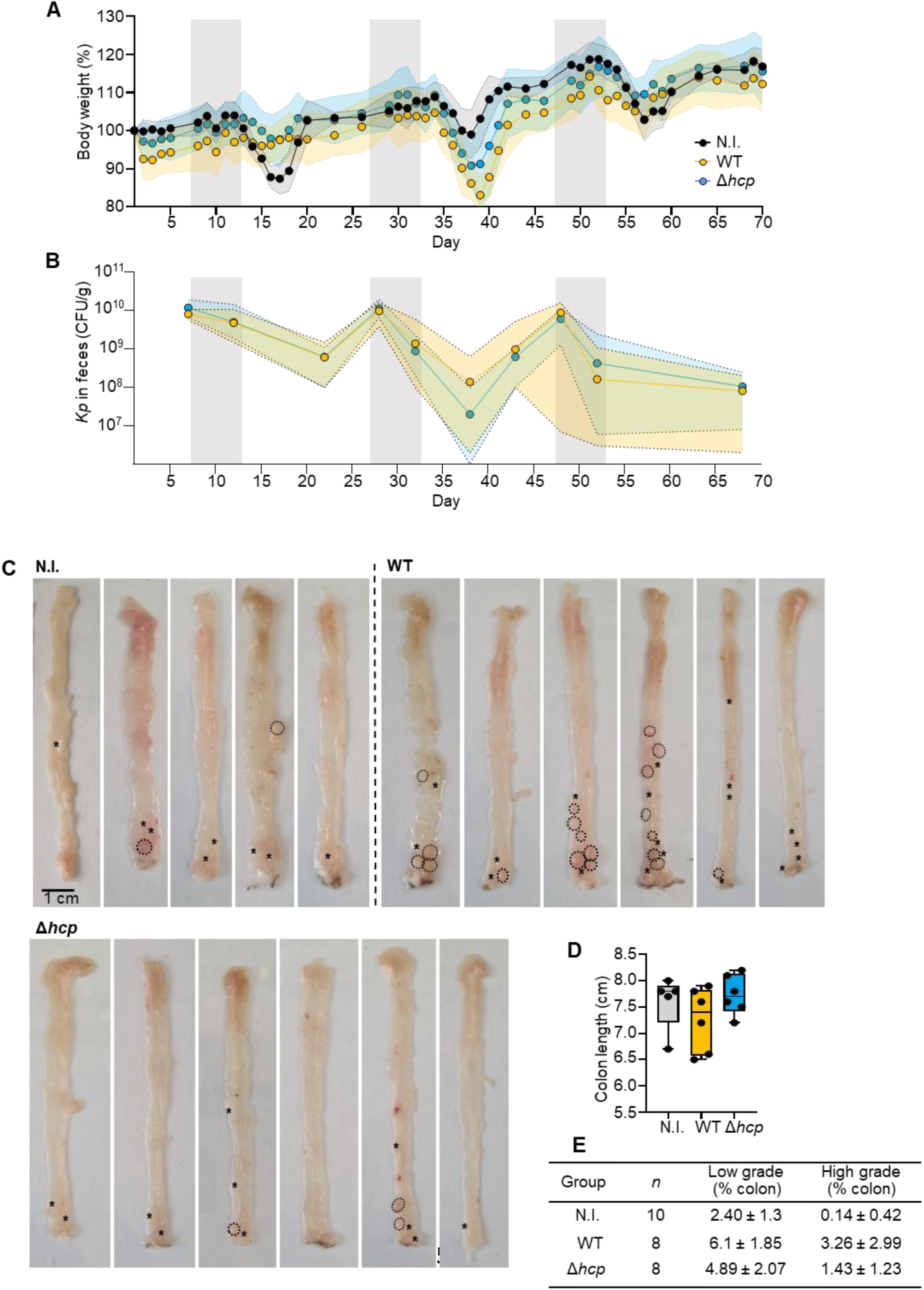
*K. pneumoniae* colonization and tumor development during AOM/DSS-induced colorectal cancer, related to Figure 6. (A) Body weight changes of mice throughout the AOM/DSS experiment. (B) *K. pneumoniae* colonization assessed by CFU counts in feces. (C) Macroscopic images of colons at sacrifice. Small tumors (<4 mm³) are indicated by stars; large tumors (>4 mm³) are outlined with dashed lines. (D) Colon length measured at the experimental endpoint. (E) Quantification of low-grade and high-grade dysplasia across experimental conditions.

